# Higher-order interactions shape microbial interactions as microbial community complexity increases

**DOI:** 10.1101/2022.05.19.492721

**Authors:** Manon A. Morin, Anneliese J. Morrison, Michael J. Harms, Rachel J. Dutton

**Affiliations:** School of Biological Science, University of California San Diego, La Jolla, 92093, USA; Department of Chemistry and Biochemistry, University of Oregon, Eugene, OR, United States; Institute of Molecular Biology, University of Oregon, Eugene, OR, United States

## Abstract

Non-pairwise interactions, or higher-order interactions (HOIs), in microbial communities have been claimed to explain the emergent features in microbiomes. Yet, the re-organization of microbial interactions between pairwise cultures and larger communities remains largely unexplored from a molecular perspective but is central to our understanding and further manipulation of microbial communities. Here, we used a bottom-up approach to investigate microbial interaction mechanisms from pairwise cultures up to 4-species communities from a simple microbiome (*Hafnia alvei, Geotrichum candidum, Pencillium camemberti* and *Escherichia coli*). Specifically, we characterized the interaction landscape for each species combination involving *E. coli* by identifying *E. coli’s* interaction-associated genes using an RB-TnSeq-based interaction assay. We observed a deep reorganization of the interaction-associated genes, with very few 2-species interactions conserved all the way up to a 4-species community and the emergence of multiple HOIs. We further used a quantitative genetics strategy to decipher how 2-species interactions were quantitatively conserved in higher community compositions. Epistasis-based analysis revealed that, of the interactions that are conserved at all levels of complexity, 82% follow an additive pattern. Altogether, we demonstrate the complex architecture of microbial interactions even within a simple microbiome, and provide a mechanistic and molecular explanation of HOIs.

## Introduction

Microbiomes are multidimensional systems containing up to thousands of interacting species. In part due to this complexity, a common strategy to investigate their biology has been to use bottom-up or reductionist approaches. Using *in vitro* microbial communities or monocultures, the features of microbial communities are measured in low-dimensional settings, with the underlying objective to understand whether high-dimension phenotypes can be predicted from lower-level observations. The lack of predictability of microbial community features from monoculture observations has demonstrated the critical importance of microbial interactions in shaping microbial communities. For instance, community assembly^1, 2^, community function^3^, community resistance to invasion^4^ or its effects on its host^5^ are highly different from the simple combination of individual species effects. Some of these studies have also shown that, while 2-species, or pairwise culture, observations could partially predict what happens in higher communities^1,2,6^, context-specific phenotypes emerge as the microbial community becomes more complex, limiting the use of pairwise information to obtain a descriptive picture of larger communities. For instance, the function (amylolytic activity) of a soil-associated microbial community of 6 species has been shown to differ from the simple linear combination (*i.e*. the addition) of the observed function of pairwise cultures^3^. Similarly, 13 to 43% of measured *Drosophila melanogaster* traits (lifespan, reproduction and development) in flies carrying a microbiome of 5 species were not predictable from the associated traits from flies inoculated with pairwise combinations of the same 5 species^5^. In the zebrafish gut microbiome, while strong negative pairwise interactions were measured, the assembly of a more complex microbiome of 5 species strikingly diverged from any assembly prediction based on pairwise interactions, and all 5 species actually co-occur in the final microbiome^7^.

Reorganization of the interaction profile with varying community complexity and the presence of higher-order interactions (HOIs) are the most likely explanation for the lack of predictability of complex community phenotypes from pairwise observations. In ecology, HOIs are described as key to stabilizing communities and promoting biodiversity in biological ecosystems ^8–10^ They are commonly described as the modification of pairwise interactions when another species is introduced, or any interaction that cannot be described by a pairwise model ^11,12^. For instance, while the bacterium *Escherichia coli* can successfully invade cultures of *Chlamydomonas reinhardtii* as well as cultures of *Tetrahymena thermophila*, it is unable to invade a coculture of the alga and the ciliate. This is due to *C. reinhardtii* inhibition of *E. coli* aggregation specifically in the presence of *T. thermophila*, which renders the bacterium vulnerable to predation^13^. Changes in microbial interactions and HOIs are undoubtedly a significant feature of microbiome ecology and functioning. Thus, investigating HOIs and deciphering how microbial interactions are rearranged when microbial systems increase in complexity are essential to our understanding of microbiomes and our ability to manipulate them.

In this work, we investigate how interactions are reorganized when the complexity of a microbial community increases from two to four species. We use a simplified cheese rind microbiome composed of the two gamma-proteobacteria *E. coli* and *Hafnia alvei*, the yeast *Geotrichum candidum* and the filamentous fungus *Penicillium camemberti*. Our previous work comparing the pairwise interaction patterns of this model microbiome to the interaction pattern in the full community highlighted the prevalence of HOIs and the lack of conservation of pairwise interactions^14^. However, we couldn’t precisely resolve the origin of these HOIs or specific rules underlying the emergence of these HOIs. For instance, we couldn’t determine whether 2-species interactions are alleviated by the introduction of a specific species or by the introduction of all the other species. Similarly, we couldn’t conclude whether community-specific interactions are actually specific to the community or whether they arise at an intermediate level and are maintained in the whole community. Relying on the ability to deconstruct and reconstruct this model system and on the RBTnSeq-based interaction assay we have previously optimized to compare gene fitness values across multiple conditions^15^, we aim to identify quantitative changes in gene fitness values for *E. coli* in different interactive conditions to identify the genetic basis of interactions at every level of community complexity.

Comparing interaction-associated genes across conditions, we can identify pairwise interaction-associated genes that are also observed at higher levels of complexity as well as genes associated with HOIs. Here, HOIs are defined as interaction-associated genes that emerge in the 3 or 4 species cultures and 2-species interaction-associated genes that are no longer observed in the 3 or 4 species cultures. Analysis of the genes associated with HOIs and their functions allowed us to further characterize the deep reorganization of the interaction landscape and highlights that it is mostly associated with the reprogramming of metabolic interactions and the introduction of a fungal partner. We further focus on the genetic basis of 2-species interactions that are conserved in higher levels of complexity to elucidate principles behind interaction conservation. We use an epistasis and quantitative genomics approach^16,17^ to understand whether interactions that are conserved follow a linear, or additive, pattern. The evaluation of interaction effects as quantitative traits allows us to define another form of HOIs as cases in which there is a lack of additivity in interactions that are conserved from simpler to more complex community composition. This form of HOIs is consistent with a more quantitative definition of HOIs, highly similar to the definition of epistasis in population genetics^18^, that identify HOIs (or epistasis) as any deviation, for a given quantitative trait, from the prediction of a linear model where only pairwise interactions are included. Carrying out this analysis, we observe that 82% of the conserved interactions follow an additive pattern of conservation from 2-species to 4-species, and that 18% of the conserved 2-species interaction is associated with non-linear models of conservation.

Overall, our work provides a unique illustration of the highly complex reorganization of interaction mechanisms when microbial community complexity changes. This provides a mechanistic explanation of HOIs in microbial communities that is essential for the further global understanding of microbial communities.

## Results

### Sets of interaction-associated genes change across interactive conditions

To investigate how microbial interactions are reorganized in a microbial community with increasing complexity, we reconstructed *in vitro* a modified bloomy rind cheese-associated microbiome on Cheese Curd Agar plates (CCA plates) as described in our previous work^14^. The original community is composed of the gamma-proteobacterium *H. alvei*, the yeast *G. candidum* and the mold *P. camemberti*. Using a barcoded transposon library of the model bacterium *E. coli* as a probe to identify interactions, we investigated microbial interactions in 2-species cultures (*E. coli* + 1 community member), in 3-species cultures (*E. coli* + 2 community members) and in 4-species cultures (or whole community: *E. coli* + 3 community members) (Figure 1A). Quantification of species’ final CFUs after 3 days of growth highlighted consistent growth for *H. alvei* and *G. candidum* independent of the culture condition and slightly reduced growth for *E. coli* in interactive conditions compared to growth alone except for following growth with *P. camemberti* (Supplementary Figure 1). Although we were unable to quantify spores of *P. camemberti* after three days, growth of *P. camemberti* was visually evident in all of the expected samples.

**Figure 1:**
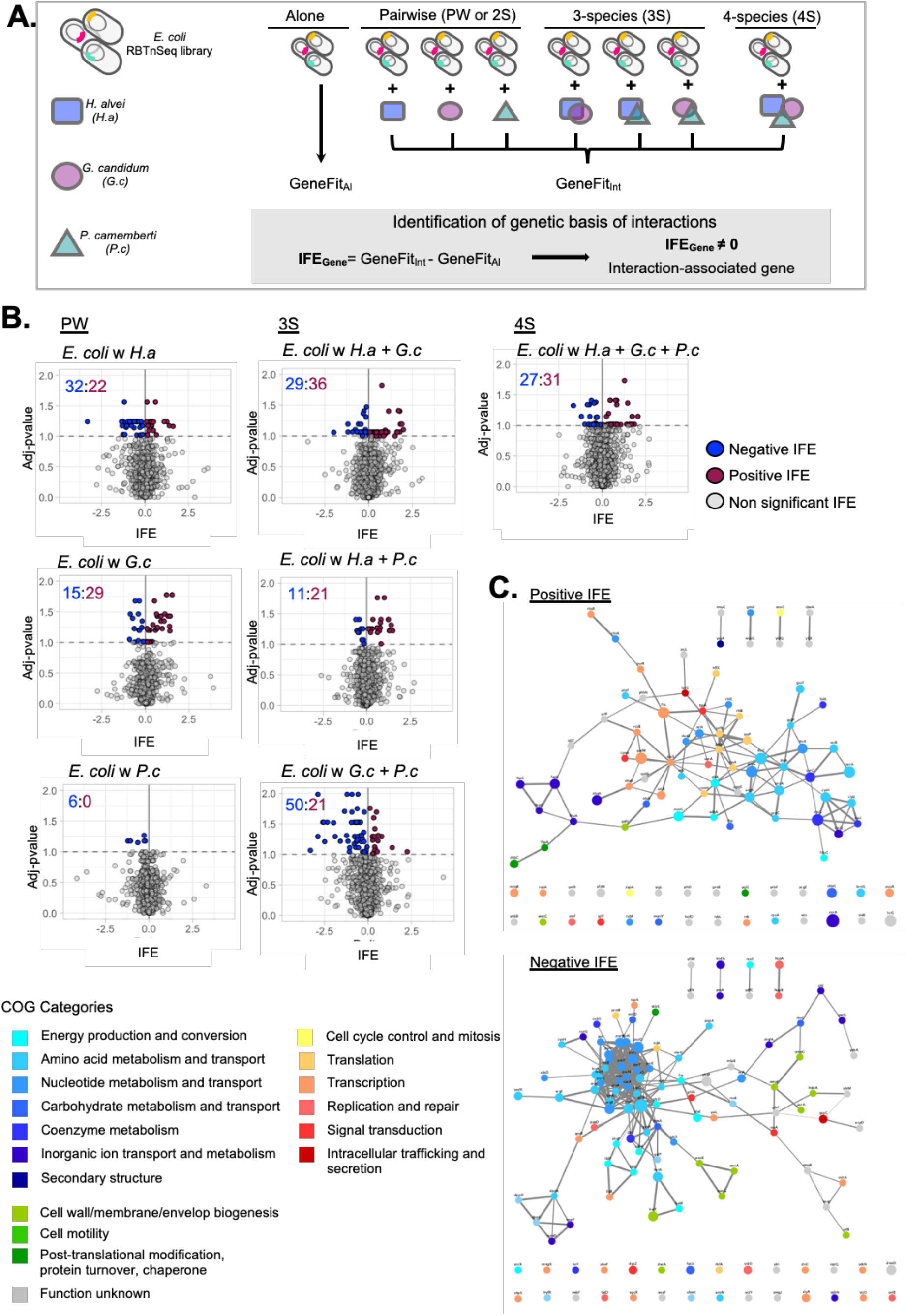
Changes of *E. coli’s* interaction-associated genes in 2-species, 3-species and 4-species cultures. **A.** Experimental design for the identification of interaction-associated genes in 7 interactive conditions from the Brie community. The *E. coli* RBTnSeq Keio_ML9 (Wetmore *et al*., 2015) is either grown alone or in 2, 3 or 4 species cultures to calculate *E. coli* gene fitness in each condition (in triplicate). Interaction fitness effect (IFE) is calculated for each gene in each interactive culture as the difference of the gene fitness in the interactive condition and in growth alone. IFE that are significantly different from 0 (two-sided t-test, Benjamini-Hochberg correction for multiple comparisons) highlight the genes associated with interaction in an interactive condition. **B.** Volcanoplots of IFEs calculated for each interactive condition. Adjusted p-values lower than 0.1 highlight significant IFEs. Negative IFEs (blue) identify negative interactions and positive IFE (red) identify positive interactions. Numbers on each plot indicate the number of negative (blue) or positive (red) IFEs. **C**. Functional analysis of the interaction-associated genes (significant IFEs). Interaction-associated genes have been separated into two groups: negative IFE and positive IFE. For each group, we represent the STRING network of the genes (Nodes). Edges connecting the genes represent both functional and physical protein association and the thickness of the edges indicates the strength of data support (minimum required interaction score: 0.4 – medium confidence). Nodes are colored based on their COG annotation and the size of each node is proportional to the number of interactive conditions in which that given gene has been found associated with a significant IFE. Higher resolution of the networks with apparent gene names are found in Supplementary Figures 2 and 3.

Previously, we developed an assay and a pipeline to identify microbial genes associated with interactions by adapting the original RB-TnSeq approach^19^ to allow for consistent implementation of biological replicates as well as for direct quantitative comparison of fitness values between different culture conditions^15^. More specifically, the original RBTnSeq assay relies on the use of a pooled library of randomly barcoded transposon mutants of a given microorganism (RBTnSeq library)^19^. Measuring the variation of the abundance of each transposon mutant before and after growth, the pipeline allows the calculation of a fitness value for each mutant. A negative fitness indicates decreased growth of the mutant relative to a wild type strain, whereas a positive fitness value indicates increased growth in the studied condition. Then, to identify genes associated with interactions, we measure and compare gene fitness across the different studied conditions, for example, comparing growth alone to growth in the presence of another species. Then, any gene whose fitness values significantly change between such conditions is identified as an interaction-associated gene.

In this work, we used the *E. coli* RBTnSeq Keio_ML9 library^19^ and grew it for 3 days alone or in the seven different interactive conditions studied here (Figure 1A). For each interactive condition, we calculated the Interaction Fitness Effect (IFE) for 3699 *E. coli* genes as the difference between the gene fitness in the studied interactive condition and the gene fitness in growth alone (Supplementary Data 1). To identify genes associated with interactions, we then tested for all the IFEs that are significantly different from 0 (adjusted p-value ≤ 0.1; two-sided t-test and Benjamini-Hochberg correction for multiple comparison^20^). Negative IFE occurs when gene fitness decreases in the interactive condition, and positive IFE occurs when gene fitness improves in the interactive condition. Here, we identified between 6 (with *P. camemberti*) and 71 (with *H. alvei* + *P. camemberti*) significant IFEs per condition (Figure 1B). Both negative IFEs and positive IFEs were found in each interactive condition except for the 2-species culture with *P. camemberti*, where only negative interactions were identified. A total of 330 significant IFEs associated with 218 unique genes were identified (as the same gene can have a significant IFE in multiple conditions) including 125 genes associated with negative IFE and 120 genes associated with positive IFE (Supplementary Figures 2 and 3).

To gain insight into the interaction mechanisms among microbes, we next analyzed the functions associated with IFEs. Here, the vast majority of genes associated with significant IFEs are part of an interaction network, highlighting the presence of genes with connected functions and from similar pathways (Figure 1C). A significant fraction of the genes associated with a negative IFE are part of amino acid biosynthesis and transport (17% - Figure 1C and Supplementary Figures 2 and 4), and more specifically with histidine, tryptophan and arginine biosynthesis. This points to competition for these nutrients between *E. coli* and the other species. Another large set of genes is related to nucleotide metabolism and transport (14% - Figure 1C and Supplementary Figures 2 and 5), highlighting competitive interactions for nucleotides and/or their precursors. The majority of these genes relate to purine nucleotides and more specifically to the initial steps of their *de novo* biosynthesis associated with the biosynthesis of 5-aminoimidazole monophosphate (IMP) ribonucleotide. Of the genes with a positive IFE, 15% are related to amino acid biosynthesis and transport (Figure 1C and Supplementary Figures 3 and 4), suggesting cross feeding of amino acids between *E. coli* and the other species. More specifically, this includes phosphoserine, serine, homoserine, threonine, proline and arginine. The presence of amino acid biosynthetic genes among both negative and positive IFEs indicate that trophic interactions (competition versus cross-feeding) depend on the type of amino-acid and/or the species interacting with *E. coli*. For both negative and positive IFEs, numerous genes were annotated as transcriptional regulators (Figure 1C and Supplementary Figures 2 and 3) emphasizing the importance of transcriptional reprogramming in response to interactions. These transcriptional regulators include metabolism regulators as well as regulators of growth, cell cycle and response to stress. Finally, these interaction-associated genes and these interaction mechanisms are consistent with previous findings in this microbiome^14^ as well as in a study of bacterial-fungal interactions involving *E. coli* and cheese rind isolated fungal species^15^.

### Introduction of a third-interacting species deeply reshapes microbial interactions

The differences in the number and sign of significant IFEs observed among the different interactive conditions, with different numbers of interaction species, suggest that the number and type of interacting partners influence interaction mechanisms. To characterize how the interactions are reorganized with community complexity, we then investigated if and how the genetic basis of interactions changes when the number of interacting partners increases by comparing the genes associated with significant IFE in 2-species cultures, in 3-species cultures and then in 4-species cultures.

First, we have identified 104 IFEs associated with 98 genes in 2-species cultures as well as 168 IFEs associated with 136 unique genes in 3-species conditions (Supplementary Figure 6 and Supplementary Data 2). Comparing these gene sets, we can identify how the interaction-associated genes change when a third-species is added to a 2-species culture. We identified 45 2-species interaction-associated genes maintained in at least one 3-species condition (maintained interaction-genes), 55 2-species interaction-associated genes no longer associated with interaction in any 3-species condition (dropped interaction-genes) and 100 3-species interaction-associated that aren’t associated with any 2-species interaction-associated genes (emergent interaction-genes) (Figure 2A, Supplementary Figure 6 and Supplementary Data 3). Both dropped and emerging interaction-associated genes represent 3-species HOIs; the third species either removes an existing interaction or brings about a new one.

**Figure 2:**
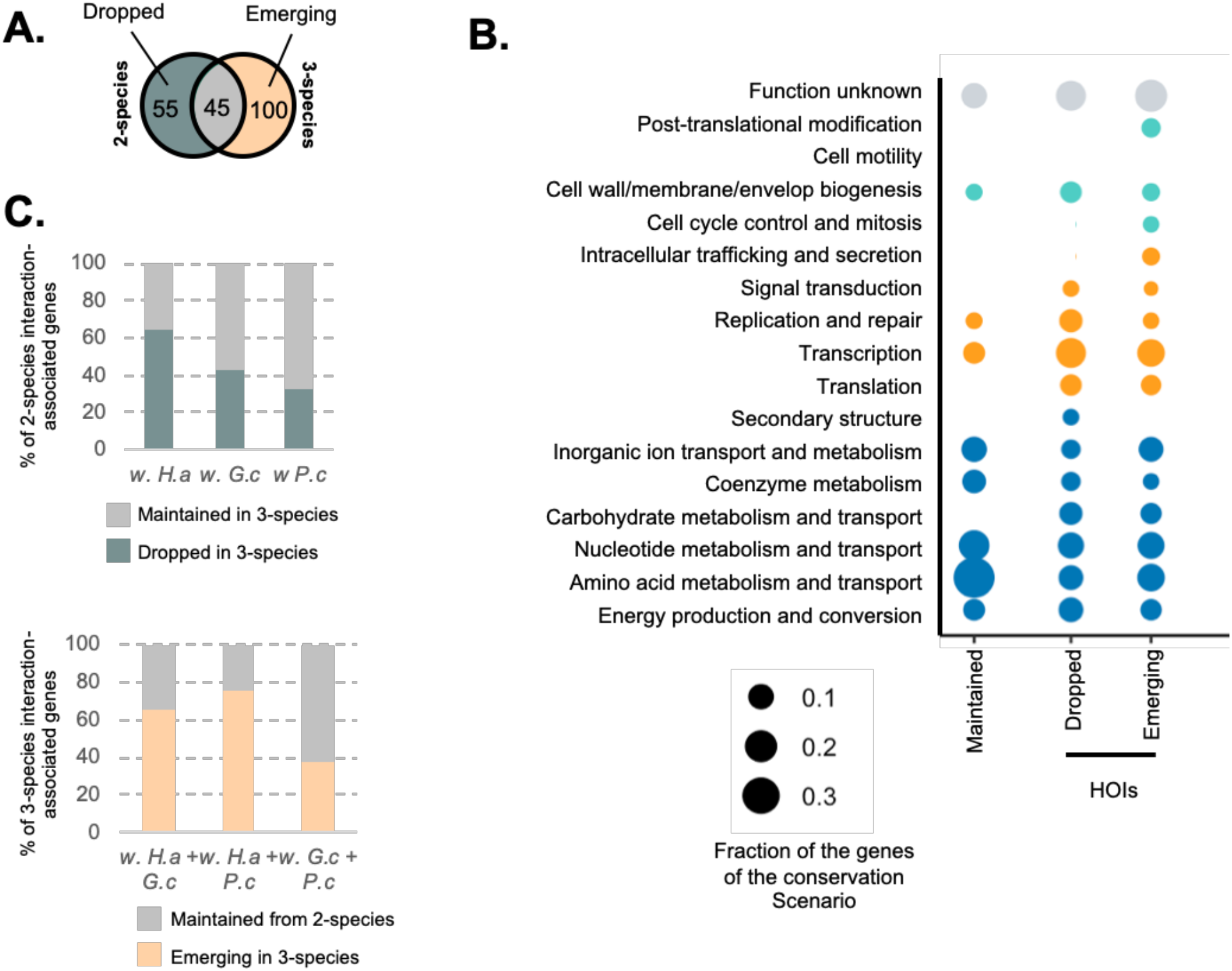
Comparison of the genetic basis of interaction for 2-species and 3-species conditions. **A.**Venn Diagram of 2-species and 3-species interaction-gene sets. This Venn Diagram identifies 2-species interaction-genes that are dropped when a third species is introduced (Left side; Dropped interaction-genes = any 2-species gene that are not found in any 3-species condition), 2-species interaction-genes that are maintained in at least one associated 3-species condition (Intersection; Maintained interaction-genes) and interaction-genes that are specific to 3-species condition (Right side; Emerging interaction-genes). **B.** Functional analysis of the dropped, maintained and emerging interaction-genes from 2-species to 3-species. Each dot represents the fraction of genes of the studied gene set associated with a given COG category (Number of genes found in the category / Total number of genes in the gene set). The color of the dots indicates the general COG group of the COG category: Teal: Metabolism; Blue: Information storage and processing; Orange: Cellular Processes and Signaling; Grey: Unknown or no COG category. **C.** Species-level analysis of 3-species HOIs: for each 2-species condition, we measure the fraction of interaction-genes that are dropped in associated 3-species cultures (Dropped in 3-species) or maintained in at least one of the 3-species cultures (Maintained in 3-species); for each 3-species condition, we measure the fraction of interaction-genes that have been conserved from at least one associated 2-species condition (Maintained from 2-species) or that are emerging with 3-species (Emerging in 3-species).

We further carried out functional analysis of maintained, dropped and emerging interaction-genes to elucidate whether maintained and HOIs interaction-genes would be associated with specific functions and thus interaction mechanisms (Figure 2B). For each set of genes, we calculated the fraction of genes of that set associated with a given COG ontology category. Metabolism and transport is the most observed COG group (Figure 2B - teal dots). For maintained interaction-genes, this indicates that some trophic interactions can be maintained from 2-species to 3-species conditions. For instance, serine biosynthetic genes *serA, serB* and *serC* as well as threonine biosynthetic genes *thrA, thrB* and *thrC* are associated with positive IFEs in the 2-species condition with *G. candidum* as well as in the 3-species conditions involving *G. candidum* (Supplementary Figure 4). This suggests that, (i) *G. candidum* facilitates serine and threonine cross feeding and (ii) this cross-feeding is still observed when another species is introduced. However, metabolism-related genes identified among the dropped and emerging interaction-agenes indicate that many trophic interactions are also rearranged through HOIs. Genes associated with lactate catabolism (*lldP* and *lldD*) and lactate metabolism regulation (*lldR*) have a negative IFE in the 2-species culture with *H. alvei*, suggesting competition for lactate between *E. coli* and *H. alvei*. Yet, these genes are no longer associated with a significant IFE when at least another partner is introduced (Supplementary Figure 7). Histidine biosynthesis genes *hisA, hisB, hisD, hisH* and *hisI* have a negative IFE in the 2-species culture *with H. alvei* and sometimes in the 3 species culture with *H. alvei* + *P. camemberti*. However, the negative IFE is alleviated whenever *G. candidum* is present, suggesting that potential competition for histidine between *E. coli* and *H. alvei* is alleviated by this fungal species (Supplementary Figure 4). Also, genes related to the COG section “Information storage and processing” are mostly found among HOIs genes, suggesting a fine-tuning of specific cellular activity depending on the interacting condition. For instance, we identified many transcriptional regulators of central metabolism among the dropped genes (*rbsR* and *lldR*) and the emerging genes (*purR, puuR, gcvR* and *mngR*), highlighting again the reorganization of trophic interactions associated with HOIs. Also, many transcriptional regulators broadly associated with growth control, cell cycle and response to stress were found among the emerging interaction-genes with 3-species (*hyfR, chpS, sdiA, slyA* and *rssB*), underlining a noticeable modification of *E. coli’s* growth environment with 3-species compare to with 2-species.

Finally, we further aimed to understand whether HOIs are associated with the introduction of any specific species (Figure 2C and Supplementary Figure 8). We observe that interaction-associated genes with *H. alvei* are more likely to be dropped, as 65% of them are alleviated by the introduction of a fungal species (Figure 2C). This can be seen, for instance, with the reorganization of *E. coli* and *H. alvei* trophic interactions following the introduction of *G. candidum* (alleviation of lactate and histine competition for instance). Also, we observe that 76% of the interactions in the 3-species cultures with *H. alvei*+*P. camemberti* and 65% in the 3-species culture with *H. alvei* + *G. candidum* are emerging genes (compared to 38% of emerging interaction-associated genes in the 3-species condition with *G. candidum* + *P. camemberti*)(Figure 2C). For the 3-species with *H. alvei + P. camemberti*, they include for instance the genes associated with purine *de novo* biosynthesis (*purR, purF, purN, purE, purC*) and the genes associated with pyrimidine *de novo* biosynthesis (*pyrD, pyrF, pyrC, carA* and *ulaD*), suggesting important trophic HOIs. For the 3-species with *H. alvei + G. candidum*, emerging interaction genes include for example the transcriptional regulators *chpS, sdiA* and *slyA*, indicating the presence of a stress inducing environment. Together, these observations suggest that the introduction of a fungal partner may introduce multiple 3-species HOIs by both canceling existing interactions and introducing new ones.

### HOIs are prevalent in a 4-species community

To further decipher whether microbial interactions continue to change with increasing community complexity, we investigated the changes in the genetic basis of interactions going from 3-species to 4-species experiments. We identified 58 interaction-genes in the 4-species condition (*E. coli* with *H. alvei* + *G. candidum* + *P. camemberti*), compared with 145 genes associated with interactions in any 3-species conditions. Comparing these two sets of genes we identify: 26 3-species interaction-genes that are maintained in the 4-species condition (including 16 directly from 2-species interactions), 115 3-species interaction-genes that are no longer associated with interactions in the 4-species condition (dropped interaction-genes) and 32 interaction-genes that are observed solely in the 4-species condition (emerging interaction-genes) (Figure 3A, Supplementary Figure 6 and Supplementary Data 3). Both dropped and emerging interactio-ngenes represent 4-species HOIs. Here, HOIs are remarkably abundant when introducing a single new species and moving up from 3-species interactions to 4-species interactions. Functional analysis of maintained and HOI genes reveals the presence of many metabolism related genes in every gene set (Figure 3), suggesting that some trophic interactions can be maintained from 3-species to 4-species interactions while some other trophic interactions are rearranged with HOIs. For instance, most of the genes of the initial steps of *de novo* purine biosynthesis have been found to have a negative IFE in the 3 species condition with *H. alvei* + *P. camemberti* (*purC, purE, purF, purL* and *purN*) as well as in the pairwise condition with *H. alvei* for *purH* and *purK* (Supplementary Figure 5), suggesting competition for purine initial precursor IMP in these conditions. Yet, the introduction of the yeast *G. candidum* as a fourth species cancels the negative IFE value, suggesting that the competition is no longer happening in its presence. Altogether, the observation of noticeable trophic HOIs moving up from 2 to 3 species and then from 3 to 4-species interaction highlights a consistent reorganization of trophic interactions along with community complexity. Also, genes related to Cell wall/membrane/envelope biogenesis are found abundantly among the 4-species emerging genes (Figure 3B) and they represent the largest functional fraction of this gene set. These genes have a negative IFE and are related to Enterobacterial Common Antigen (ECA) biosynthetic processes (*wecG, wecB* and *wecA*) (Supplementary Figure 9). While the roles of ECA can be multiple but are not well defined^21^, they have been shown to be important for response to different toxic stress, suggesting the development of a specific stress in the presence of the four species.

**Figure 3:**
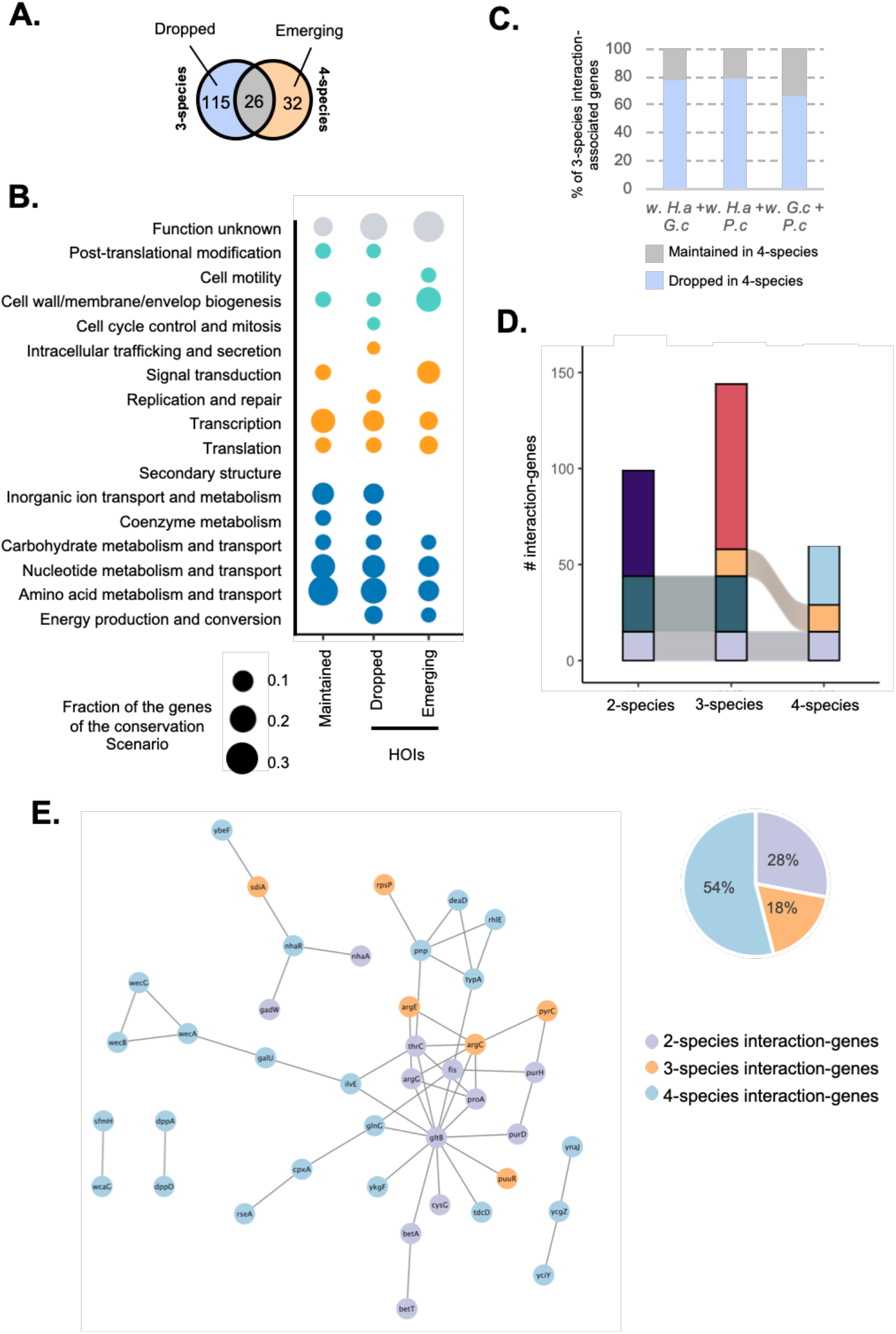
Organization of the interactions in the 4-species community. **A.** Venn Diagram of 3-species and 4-species interaction-gene sets. This Venn Diagram identifies 3-species interaction-genes that are dropped when a fourth species is introduced (Left side; Dropped interaction-genes = any 3-species gene that are not found in the 4-species condition), 3-species interaction-genes that are maintained in the 4-species condition (Intersection; Maintained interaction-genes) and interaction-genes that are specific to 4-species condition (Right side; Emerging interaction-genes). **B.** Functional analysis of the dropped, maintained and emerging interaction-genes from 3-species to 3-species. Each dot represents the fraction of genes of the studied gene set associated with a given COG category (Number of genes found in the category / Total number of genes in the gene set). The color of the dots indicates the general COG group of the COG category: Teal: Metabolism; Blue: Information storage and processing; Orange: Cellular Processes and Signaling; Grey: Unknown or no COG category. **C.** Species-level analysis of 4-species HOIs: for each 3-species cultures we measure the fraction of interaction-genes that is conserved in the 4-species culture (Maintained in 4-species) and the fraction of interaction-genes that has been dropped (Dropped in 4-species). **D.** Alluvial plots of the interaction genes across community complexity levels. **E**. STRING network of the 4-species interaction genes (Nodes). Edges connecting the genes represent both functional and physical protein association and the thickness of the edges indicates the strength of data support (minimum required interaction score: 0.4 – medium confidence). Nodes are colored based on the level of community complexity the genes are conserved from.

As for the 2 to 3 species comparison, we investigated whether the introduction of a specific fourth species would be most likely associated with HOIs. The 3-species culture that appears to be the least affected by the introduction of a fourth member is with *G. candidum* + *P. camemberti* where 34% of the observed interactions are still conserved in the 4-species condition after the introduction of *H. alvei* (versus 22% for with *H. alvei* + *G. candidum* when *P. camemberti* is added and 21% for with *H. alvei* + *P. camemberti* when *G. candidum* is added) (Figure 3C and Supplementary Figure 10). Together, these observations suggest that, again, the introduction of a fungal partner may introduce multiple 4-species HOIs.

Finally, by increasing the number of interacting species in our system and investigating interaction gene maintenance and modification with every increment of community complexity, we are able to build our understanding of the architecture of interactions in a microbial community. Altogether, we have observed a total of 218 individual genes associated with interactions in any experiment. Only 16 of them (7%) were conserved across all levels of community complexity (Figure 3D). Starting from 2-species interaction genes, 48% of them were maintained with 3-species and only 15% (16 out of 104) were still maintained with 4-species. Thus, we demonstrate here a progressive loss and replacement of 2-species interactions as community complexity increases and the prevalent apparition of HOIs. Tracking back the origins of the genetic basis of interactions in the 4-species experiment that represents the full community of our model, we identify that 28% of the full community interactions can be traced back to 2-species interactions, 18% are from 3-species interaction and 54% are specific to the 4-species interaction (Figure 3D and 3E). Most of the maintained interaction-genes from 2-species as well as from 3-species are associated with metabolism (Figure 3D and Supplementary Figure 11) while Signal transduction and cell membrane biosynthesis genes are most abundant among the 4-species interaction-genes as previously mentioned. To conclude, this shows that the genetic basis of interactions and thus the sets of microbial interaction are deeply reprogrammed at every level of community complexity and illustrates the prevalence of higher order interactions (HOIs) even in simple communities.

### The majority of maintained 2-species interaction-genes in the 4-species culture follows an additive conservation behavior

While HOIs are abundant in the 4-species condition, our data yet suggest that up to 28% of the interactions are maintained from 2-species interactions. However, we don’t know whether and how 2-species interactions are quantitatively affected by the introduction of other species and whether they would follow specific quantitative models of conservation. For instance, we can wonder how the strength of a given 2-species interaction is modified by the introduction of one or two other species, or how two 2-species interactions associated with the same gene will combine when all the species are present. In other words, can we treat species interactions as additive when we add multiple species? Such information would generate a deeper mechanistic understanding of the architecture of microbial interactions while allowing us to potentially predict some whole community interactions from 2-species interactions. Here, two main hypothetical scenarios can be anticipated. First, the conservation of 2-species interactions follows a linear or additive behavior, where the introduction of other species either doesn’t affect the strength of the conserved 2-species interaction or two similar 2-species interactions combine additively. The second scenario identifies non-linear or non-additive conservation of 2-species interactions, where the strength of the conserved 2-species interaction is modified by the introduction of other species or two similar 2-species interactions are not additive. The second scenario would encompass for instance synergistic effects or inhibitory effects following the introduction of more species. We next use an epistasis and quantitative genomics approach to understand whether interactions that are conserved follow a linear, or additive, pattern. For the 16 genes that are associated with interaction in 2-species cultures, in associated 3-species cultures and in the 4-species condition, we use epistasis analysis to test the linear behavior of IFE when the number of interacting species increases, as IFEs are quantitative traits related to the interaction strength. In multi-dimensional systems, an epistasis analysis quantifies the additive (or linear) behavior of conserved quantitative traits. In quantitative genetics, for instance, epistasis measures the quantitative difference in the effects of mutations introduced individually versus together^18,22,23^. Using a similar rationale, we can use IFEs as a quantitative proxy for interaction strength and test whether the IFEs of the maintained interaction genes in 3-species and in 4-species conditions result from the linear combination of associated 2-species IFEs (Figure 4A). Nonlinear combination, or non-additivity of 2-species IFEs in higher community level also highlights higher-order interactions.

**Figure 4:**
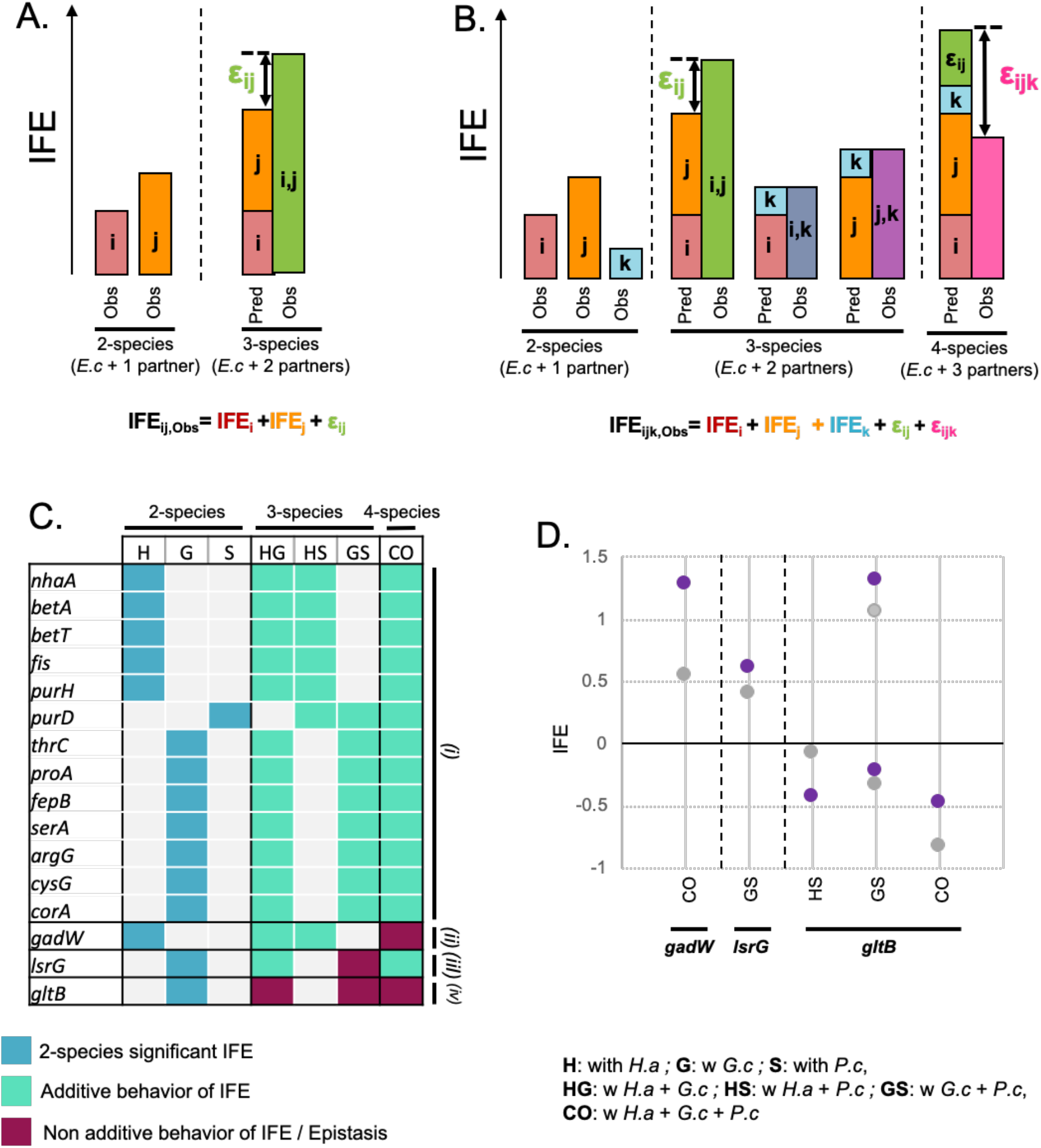
Quantitative analysis of IFE conservation for the interaction associated genes conserved from 2-species to 4-species conditions. **A.** Schematized quantitative epistasis/non-linearity measured in 3-species conditions (with partner i and j). Epistasis (ε_ij_) is the difference between the individual IFE of partner i and partner j (red and orange bars) versus placing them together (green). Mathematically, we need three terms (IFE_i_, IFE_j_, and ε_ij_) to reproduce the observed IFE for the 3-species condition. **B.** This analysis can be extended to higher levels of community complexity: 4-species (*E. coli* with 3-partners i, j, and k). The model first accounts for epistasis between i/j, i/k, and j/k. In this example, i and j exhibit epistasis; i/k and j/k are additive (dark blue and purple). The predicted IFE for the 4-species community is the sum of the individual 2-species effects (red, orange, light blue) and the 3-species epistatic terms (green). The 4-species epistatic coefficient is the difference between this low-order prediction and the observed IFE for the i,j,k community (pink). **C.** Conservation profiles of the 16 2-species interaction-associated genes conserved up to 4-species. 2-species conditions: a colored square indicatea the 2-species condition(s) in which the interaction-associated gene was identified; a grey square indicates non-significant 2-species IFEs. 3-species conditions: a teal square indicates that the associated IFE is associated with additive behavior from associated 2-species IFE (no εij epistatic coefficient), a red square indicates that the associated IFE display non-additivity from 2-species IFE and thus epistasis, a grey square corresponds to a 3-species condition that is not associated with significant 2-species IFE (no epistasis analysis performed); 4-species condition: a teal square indicates that the associated IFE is associated with additive behavior (no ε_ijk_ epistatic coefficient), a red square indicates that the associated IFE is associated with non-additivity from lower-order IFE. **D.** Comparison of the observed and predicted IFE for the genes and condition associated with 3-species and 4-species non-additive IFE.

We adapted the pipeline Epistasis^17^, originally designed for quantitative genetics investigation. We implemented the linear model with the gene fitness values for growth alone, for each of the 2-species conditions, for each of the 3-species cultures and for the 4-species condition. For each gene, the software finds the simplest mathematical model that reproduces the observed IFEs across all levels of community complexity. In the simplest case, the model will have a term describing the effects for adding each species individually to the *E. coli* alone culture; that term corresponds to the 2-species IFE. Then, if the IFE for two *E. coli’s* partners combined (3-species IFE) differs from the sum of their individual effects (corresponding 2-species IFE), the software adds a term capturing this epistasis (Figure 4A). Here, we call that term 3-species epistatic coefficient or ε_i,j_ Finally, if the IFE for the combined community (*E. coli* plus all three species; 4-species condition) differs from the prediction based on the 2-species and 3-species terms, the software will add a high-order interaction term to the model (Figure 4B). Here, we name that term 4-species epistatic coefficient or ε_ijk_.

We performed this analysis on the 16 genes that are associated with interactions at every level of community complexity. To identify real additive behavior of IFE from non-additivity, we screen for 3-species epistatic coefficients and 4-species epistatic coefficients that are significantly different from 0 (adjusted p-value ≤ 0.01, correction for multiple testing Benjamin-Hochberg). We found that 13 genes behaved additively from 2-species to 4-species culture, with no epistatic contributions in the 3-species conditions nor in the 4-species condition (Figure 4C, *(i)*). One gene (*gadW*) exhibited nonlinear conservation of IFE only in the 4-species condition, but additive IFE conservation from 2-species to 3-species (Figure 4C, *(ii)*). Another gene (*lsrG*) showed epistasis in one 3-species condition but no epistasis in the 4-species condition (Figure 4C, *(iii))* Finally, one gene (*gltB*) displayed both non-additivity in 3-species and 4-species conditions (Figure 4C, *(iv)*).If we look more closely at the additive genes, we find genes *(betA, betT, purD* and *purH*) are associated with the conservation of negative IFEs (Supplementary Figure 12). While *betA* and *betT* are associated with choline transport (*betT*) and glycine betaine biosynthesis from choline (*betA*)^24^, *purD* and *purH* are associated with *de novo* purine biosynthesis^25^. This suggests that requirements for glycine betaine biosynthesis from choline and for purine biosynthesis caused by microbial interactions, possibly due to competition for the nutrients used as precursors, are additively conserved from individual 2-species interactions requirements. Also, 5 genes associated with amino acid biosynthesis (*serA, thrC, cysG, argG* and *proA*) are associated with the additive conservation of positive IFE (Supplementary Figure 12), suggesting that cross feeding can be additive when the community complexity increases. Altogether, this highlights the existence of 2-species interactions, including trophic ones, conserved in an additive fashion in the highest-level of complexity.

This leaves 3 genes (18%) of the maintained 2-species interaction-associated genes that are associated with non-additive behavior, and thus HOIs, at at least one higher level of community complexity (Figure 4C - *(ii), (iii)* and *(iv)*). The gene *gadW* is associated with non-additivity at the 4-species level, suggesting that while IFEs are additive in 3-species cultures, the introduction of a fourth species introduces HOI. Moreover, the observed 4-species IFE is greater than the IFE predicted by a linear model (Figure 4D), highlighting a potential synergistic effect when the 4 species are together. The gene *lsrg* is associated with non-additivity only at the 3-species culture w *G.c* + *P.c*. More specifically, this indicates that HOI arise when these 2 fungal species are interacting together with *E. coli*, but that no more HOI emerge when *H. alvei* is introduced (ie, the 4-species IFE can be predicted by the linear combination of the lower levels IFEs). As the observed IFE for the 3-species condition w *G.c* + *P.c* is greater than the predicted IFE (Figure 4C), this suggests a synergistic effect between the 2 fungal species. Finally, the gene *gltB* is associated with non-additivity at both the 3-species and 4-species levels. For this gene, the conservation of IFE is never associated with an additive model. Here, the observed 4-species IFE is not as negative as it would be as the result of the linear combination of the associated lower IFE (Figure 4D), suggesting the existence of a possible IFE threshold, or plateau effect. Altogether, this indicates that maintained 2-species-interactions can follow nonlinear behaviors that could involve synergistic effects, inhibitory effects or constraints.

## Discussion

Interactions between microbes are responsible for the specific and multiple phenotypes observed in microbial communities compared to monocultures. Knowledge of microbial interactions could be the key to controlling microbial communities, but deciphering these interactions is challenging in complex microbiomes. Moreover, pairwise culture interactions are often insufficient to predict what happens in more complex microbiomes, suggesting an important reprogramming of interactions as community complexity increases in terms of the number of species present^3,5,7,13,14^. Understanding the restructuring of these interactions in complex communities is thus essential to comprehending the biology of microbial communities. Using an *in vitro* multi-kingdom community, we performed a molecular investigation of the reorganization of interaction profiles as community complexity changed. Relying on the tractability of our model system and an RBTnSeq-based interaction assay, we tracked interactions in 2-species, 3-species and 4-species cultures. In this work, the combination of a qualitative and quantitative comparison of interaction profiles at the molecular level underlines the complex dynamics of interaction reorganization with community complexity and the existence of multiple forms of higher-order interactions.

This work offers an example of the different forms that HOIs can take in biological systems. We report multiple mechanistic HOIs as defined in ecological studies and represented here by any pairwise interaction-associated genes that are not observed in 3 and more species conditions as well as any interaction-associated genes observed in higher-levels than pairwise cultures^12^. We also report another form of HOIs, as defined in quantitative genetics, which are associated with the non-additive behavior of conserved 2-species interactions in 3 or 4 species communities^22^. Here, each HOI level refers to different biological phenomena occurring in the same biological system. Yet, they are essential and complementary to decipher the extremely convoluted architecture and dynamics of microbial interactions and microbiome biology.

As the number of interaction-associated genes strongly decreased in the 4-species culture compared to 2 or 3 species setups, our work points out a strong reduction of the interaction landscape with 4 species that is associated with the loss of many lower-level interactions and the emergence of context-specific interactions. While 43% of the 2-species interaction-associated genes are still found with 3 species, only 15% are still found with 4 species, representing less than a third (28%) of the total of 4-species interactions. This highlights the increasing dilution and replacement of original 2-species interactions. To summarize, the more complex a community gets, the more mechanistic HOIs emerge. While our work was limited to 4 species, it remains necessary to verify whether this statement will be true for more complex communities: whether more 2-species interactions will be lost and more HOIs will keep emerging at each level or whether the interaction landscape will stabilize. Indeed, Friedman *et al*., have highlighted that growth observation for 2- and 3-species combination could predict the assembly of 7- to 8-species^2^; this would suggest that interactions, or at least key interactions driving community assembly, could change less dramatically in higher complexity microbiomes.

Our molecular approach enabled us to identify that most of the reorganization of the interaction profile is associated with the reprogramming of metabolic or trophic interactions including both competition for nutrients and cross-feeding. As more species are introduced it appears that the dynamics of nutrient consumption is rearranged. For instance, we observed that some 2-species competition for amino acids and for lactate between *E. coli* and *H. alvei* can be alleviated by the introduction of *G. candidum*. While the competition for amino acids is likely alleviated by amino acid cross-feeding from *G. candidum*, as this species is known to release amino acids into the environment through digestion of proteins and peptides^26–28^, the mechanism relieving competition for lactate is unclear. Possibly, *G. candidum* provides other nutrients sources like amino acids that alleviate the need for *E. coli* to rely on lactate. Trophic interactions are described as core determinants in community assembly^29,30^ and simple rules of metabolic interdependencies and metabolic specialization can be sufficient to predict the assembly of rather complex communities^31–33^. Yet in complex systems, such as the gut, with complex nutrient composition, the important reorganization of metabolic interactions as community complexity changes could likely explain the difficulty in predicting community assembly and composition in some studies^7^. As previously suggested^34^, ecological and metabolic factors associated with the present species such as niche overlap, degree of metabolic specialization and species similarities are likely to be drivers of these metabolic HOIs. Indeed, microorganisms display incredible metabolic abilities, from nutrient usage to rapid metabolic switching, and in the presence of multiple nutrient sources and/or other microorganisms they are likely to readjust the sequence of nutrient uptake. Niche occupation and nutrient access are also two crucial aspects in the success or failure of microbial invasion^4,35^. The reorganization of metabolic interactions in different community composition could likely explain the poor predictability of invasion resistance from simple species-combinations and the emergence resistance-specific phenotype.

To some extent, our work also highlights the importance of fungal species as major actors in reshaping interaction networks. Here, more 3-species HOIs and 4-species HOIs were observed when one fungal species was added either as a third species with two bacteria or as a fourth species with another fungus and two bacteria. In microbial communities, fungi are known to impact community structure and access to nutrients through the formation of hyphal highways^36,37^, to impact community assembly through environmental modification^38–40^ as well as to impact community protection through the production of multiple secondary metabolites with antibiotic or antimicrobial properties^15,39,41^. In this specific context, fungi-associated HOIs seem to be related to their metabolic characteristics, whether through cross-feeding or competition for nutrients. Yet, the emergence of specific requirements for ECA biosynthetic genes in *E. coli* when all species are present, suggesting a potential toxic stress, may reveal other fungal context-specific properties, while it could also inform about a more precise role of the ECA in *E. coli*. We believe this work contributes, along with other recent studies, to advocate for the need to include fungal species more frequently into microbiome work studies.

Finally, to understand principles behind the conservation of pairwise interactions, we used epistasis analysis to quantify the additive or non-additive behavior of conserved 2-species interactions in 3 and 4 species conditions. Epistasis analysis offers an adaptable approach to test the linear behavior of quantitative traits in systems with increasing complexity, whether it is the effects of accumulation of mutations in a genome^18,42^, the effect of drug combinations^43^ or the dynamics of predators-prey ecosystems^44^. Recent studies in the field of microbiome research have also relied on epistasis analysis to elucidate fundamentals of microbial communities such as how the nutrient composition of the environment determines the assembly and the diversity of a microbial community^34,45^, how the functional landscape of microbial community is built^3^ or how the commensal microbiome determines the development, lifespan and reproduction of its host^5^. In this work, using the quantitative metric IFE for the strength of interactions, we used an epistatic model to characterize the additivity or non-linearity of IFE of conserved 2-species interactions when *E. coli* is growing with 2 and 3 other species (3-species and 4-species conditions respectively). We observed that most of the conserved 2-species interaction genes followed an additive model of conservation, including trophic-interaction related genes. While our study was limited to a small number of interacting species, it remains to be investigated if this linear behavior is propagated if the community complexity increases again. While we have observed both synergistic effects and limitations effects for nonlinear conservation of interaction strength, we believe that such behavior are likely to arise in more complex contexts and that additivity will eventually stop. For instance, additivity of pairwise interactions is likely to saturate and to stop at higher-levels of complexity due to environmental constraints (nutrient supply for instance) and/or the metabolic capacities of the different species as highlighted in ^34^. While this theory can apply to conserved trophic interactions, like the ones highlighted in our work, other scenarios could include molecule-driven or environment-mediated interactions such as exchange of metabolites, quorum sensing, antibiosis and modification of the physiochemical properties of the environment. While trait additivity is fairly easy to quantify in high-dimensional systems, our study, along with other work using epistasis to study microbial communities^3,5,34,45^ suggest that additivity of quantitative microbial features is not the only scenario in nature, and strongly emphasizes the need to develop mathematical approaches to accurately understand the dynamics and functionalities of microbiomes.

Overall, our work identifies the reorganization of microbial interactions along with community complexity at the molecular level. While more advanced modeling is still required, this knowledge of the mechanistic reorganization of interactions and patterns of HOIs is essential for bottom-up approaches to be able predict, from minimal information, the biology of complex microbiomes and will offer novel opportunities to design synthetic microbial communities or control natural ones.

## Methods

### Strains and media

#### Strains

The bloomy rind cheese community was reconstructed with the same strains from ^14^: *H. alvei* JB232 isolated from cheese^46^ and two industrial cheese strains: *G. candidum* (Geotrichum candidum GEO13 LYO 2D, Danisco – CHOOZITTM, Copenhagen, Denmark) and *P. camemberti* (PC SAM 3 LYO 10D, Danisco - CHOOZITTM).

#### Medium

All assays have been carried out on 10% cheese curd agar, pH7 (CCA) (10% freeze-dried Bayley Hazen Blue cheese curd (Jasper Hill Farm, VT), 3% NaCl, 0.5% xanthan gum and 1.7% agar). The pH of the CCA was buffered from 5.5 to 7 using 10M NaOH.

### Competition assay – RBTnSeq Assays

The *E. coli* barcoded transposon library Keio_ML9^19^ was used for all RBTnSeq assays on CCA during a 3-day growth in eight different culture conditions: alone, 2-species conditions (with *H. alvei;* with *G. candidum;* and with *P.camemberti*), 3-species conditions (with *H. alvei* + *G. candidum*; with *H. alvei* + *P. camemberti*; and with *G. candidum* + *P. camemberti*), and in the single 4-species condition (with *H. alvei* + *G. candidum* + *P. camemberti*) (Figure 1).

From the species and library inoculation to cell harvest after 3 days of growth, we followed the same procedure as described in^14^. We amplified the *E. coli* barcoded transposon library Keio_ML9 into 25 mL of liquid LB-kanamycin (50 mg/mL) up to an OD of 0.6-0.8. A 5mL sample of this preculture was spun down and kept at −80C as the T0 sample required for fitness calculation. The remaining cells were then washed in PBS1x-Tween0.05% and used to inoculate the competition assays with 7*10^6^ cells of the library on each 100 mm petri dish plate. When necessary, the other community members were then inoculated at the following densities: for *H. alvei:* 7*10^6^ cells; for *G. candidum*: 7*10^6^ cells; for *P. camemberti:* 7*10^5^ cells.

After 3 days, cells were harvested by flooding the plates with 1.5mL of PBS1X-Tween0.05%, gentle scraping and then transferring the resuspended cells into 1.5mL tubes. Cells were centrifuged for 3 min at RT at 10 KRPM and stored at −80C until gDNA extraction.

All assays have been performed in triplicate.

### gDNA extraction, library preparation and sequencing

gDNA from the samples of the competition assays was extracted by phenol-chloroform extraction (pH 8) as described in ^14^. For each sample, we resuspended the cells into 500 mL of buffer B (200 mM NaCl, 20 mM EDTA) and then transferred them into a 2mL screw-capped tube previously filled with 125 mL of 425-600 mm acid-washed beads and 125 mL of 150-212 mm acid-washed beads. Then, 210mL of SDS 20% and 500mL of Phenol:Chloroform (pH 8) were added to each sample before mechanical lysis by vortexing the tubes for 2 minutes at maximum speed. Tubes were then centrifuged for 3 min at 8 KRPM at 4C and 450mL of aqueous phase were recovered for each sample. 45 mL of sodium acetate 3M and 450 mL of ice-cold isopropanol were added and tubes were incubated for 10 min at −80C. Tubes were then centrifuged for 5 min at 4C at 13 KRPM, supernatant was removed and the pellet was washed in 750 mL of 70% ice-cold ethanol before being resuspended in 50mL of DNAse/RNAse free water.

For library preparation, the 98C BarSeq PCR described in ^19^ was used to amplify the barcoded region of the transposons and PCR was performed in a final volume of 50 mL: 25 mL of Q5 polymerase master mix (New England Biolab), 10 mL of GC enhancer buffer (New England Biolab), 2.5 mL of the common reverse primer (BarSeq_P1^19^) at 10 mM, 2.5 mL of a forward primer from the 96 forward primers (BarSeq_P2_ITXXX^19^) at 10 mM and 50 ng to 2 mg of gDNA. The following PCR program was used: (i) 98 °C - 4 min, (ii) 30 cycles of: 98 °C – 30 s; 55 °C – 30 s; 72 °C – 30 s, (iii) 72 °C – 5 min. After the PCR, 10 mL of each of the PCR products were pooled together to create the BarSeq library. 200 mL of the pooled library were purified using the MinElute purification kit (Qiagen) and final elution of the BarSeq library was performed in 30 mL in DNAse and RNAse free water.

The BarSeq library was quantified using Qubit dsDNA HS assay kit (Invitrogen) and then sequenced on HiSeq4000 (50 bp, single-end reads), by the IGM Genomics Center at the University of California San Diego.

### Data processing and interaction fitness effects (IFE) analysis

For each library, BarSeq reads were first processed using the Perl script BarSeqTest.pl from^19^ to obtain the count file (all.poolcount) containing the number of reads per barcode for each sample. This pipeline requires a table where each barcode is mapped to a location in the genome. The Arkin lab (Physical Biosciences Division, Lawrence Berkeley National Laboratory, Berkeley, California, USA) kindly provided the TnSeq table for the *E. coli* library. The original script used for this analysis originates from ^19^ is publicly available on https://bitbucket.org/berkeleylab/feba.

The generated all.poolcount information was then implemented into custom R scripts from^15^ to determine the average fitness scores for each gene across three RBTnSeq assay replicates for each of the eight cultures (https://github.com/DuttonLab/RB-TnSeq-Microbial-interactions). Insertion mutants that did not have a sufficient T0 count in each condition or that were not centrally inserted (10–90% of gene) were removed from the analysis and counts were then normalized using a set of five reference genes (*glgP, acnA, modE, leuA* - average of 52 strains each). Detailed explanation of the fitness calculation strategy can be found in the Readme document as well as in^15^. In each condition, to assess the possible variability between replicates, we measured the correlation between the three replicates (Supplementary Figure 13). In this set of experiments, Pearson coefficient between replicates varied from 0.79 to 0.86 suggesting low technical noise.

Gene fitness values were then compared between interactive conditions and *E. coli* growth alone conditions using two-sided *t*-tests (when the equality of variance was verified by Fisher test) and correction for multiple comparison (Benjamini–Hochberg method^20^). Comparisons associated with an adjusted *P* value lower than 10% were considered a significant interaction fitness effect.

### Epistasis model analysis

We used the Python package Epistasis^17^ to run the epistasis analysis on *E. coli’s* genes. We implemented the model with the average fitness values across the three biological replicates along with the corresponding variance, for each gene and for each of the culture conditions (Alone, w *H.a*, w. *G.c*, w *P.c*, w *H.a* + *G.c*, w *H.a* + *P.c*, w *G.c* + *P.c*, w *H.a* + *G.c* + *P.c*). The model is run for each gene individually. The ‘genotypes’ expected in the model correspond to the culture conditions and are binary coded based on the presence or not of *E. coli’s* partners *G. candidum, H. alvei* and *P. camemberti*. For instance, the Alone condition corresponds to phenotype 000, the w *G.c* condition is coded 100, the w *H.a*+*P.c* condition is coded 011, the w *H.a* + *G.c* + *P.c* is coded 111 and so on. The ‘phenotype’ implemented in the model corresponds to the average fitness value across three biological replicates. After generating the genotype-phenotype map for each gene, the model was run a first time using the ‘local’ parameter to set-up the center of the map, and returns the set of epistatic coefficients for each gene calculated as follows:

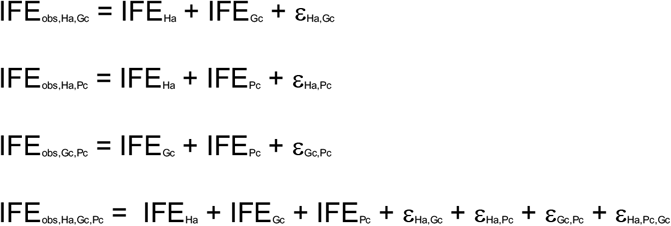

To determine whether each coefficient was different from zero, we generated 7,000 pseudoreplicates from our experimental variance from each gene and then calculated the probability each coefficient was above or below zero from this distribution. P-values were then adjusted for multiple comparison testing using Benjamini Hochberg Correction^20^. As it has been shown previously that epistatic analyses can fail if the effects of mutations combine non-linearly instead of linearly (e.g. mutational effects multiply rather than add), we used the epistasis package to look for such nonlinearity. However, no nonlinearity was observed (Supplementary Figure 14).

## Acknowledgments

The authors would like to thank the following people and groups: the Arkin Lab and the Deutschbauer Lab at UC Berkeley for the *E. coli* Keio_ML9 RB-TnSeq library; K. Jepsen at the IGM Genomics Center at the University of California, San Diego for assistance with sequencing; and all members of the Dutton Lab for constructive comments on the manuscript. This work was supported by the National Institute of Health New Innovator grant #DP2 AT010401 (R.J.D), the NIH Institutional Training grant NIH-T32GM007413 (A.J.M), National Science Foundation CAREER Award DEB-1844963 (M.J.H),

## Author Contributions

R.J.D., M.J.H. and M.A.M. conceptualized the study. M.A.M. performed the experiments. The Epistasis pipeline was written and adapted by M.J.H. and A.J.M. Data analyses were performed by M.A.M. and A.J.M. The article was written by M.A.M. and revised with input from all authors. The figures were made by M.A.M with input from all authors. The study was supervised by R.J.D. and M.J.H.

## Additional information / Competing Interest Statement

The authors declare that no financial and non-financial competing interests exist in relation to the work described.

## Supplementary material list

### Supplementary Figures

<u>Supplementary Figure 1</u>: Growth of the community species in the studied conditions

<u>Supplementary figure 2:</u> Functional analysis of interaction-associated genes with negative IFEs

<u>Supplementary figure 3</u>: Functional analysis of interaction-associated genes with positive IFEs Supplementary figure 4: IFE profiles of Amino acid biosynthesis genes identified in this study

<u>Supplementary figure 5</u>: IFE profiles of Purine biosynthesis associated genes identified in this study

<u>Supplementary Figure 6</u>: Comparison of interaction-associated genes across the different levels of community complexity

<u>Supplementary figure 7</u>: IFE profiles of lactate metabolism genes

<u>Supplementary figure 8</u>: Condition specific comparison of interaction-associated genes for 2 and 3-species conditions

<u>Supplementary figure 9</u>: IFE profiles of Enterobacterial Common Antigen (EAC) genes

<u>Supplementary figure 10</u>: Condition specific comparison of interaction-associated genes for 3 and 4-species condition

<u>Supplementary figure 11</u>: Functional network of the 4-species interaction-genes and their origin

<u>Supplementary figure 12</u>: IFE profiles of the 16 2-species interaction genes maintained up to 4-species

<u>Supplementary figure 13</u>: Pearson correlation of gene fitness across replicates

<u>Supplementary figure 14</u>: Non-linearity analysis of IFE in the Epistatis model

### Supplementary Datasets

<u>Supplementary Data 1</u>: RB-TnSeq based interaction analysis (Fitness values, Interaction Fitness Effects and associated statistics)

<u>Supplementary Data 2</u>: Interaction-associated genes at each level of community complexity

<u>Supplementary Data 3</u>: Comparison of interaction-associated genes across the different levels of community complexity

