## Supplementary Figures for "Higher-order interactions shape microbial interactions as microbial community complexity increases"

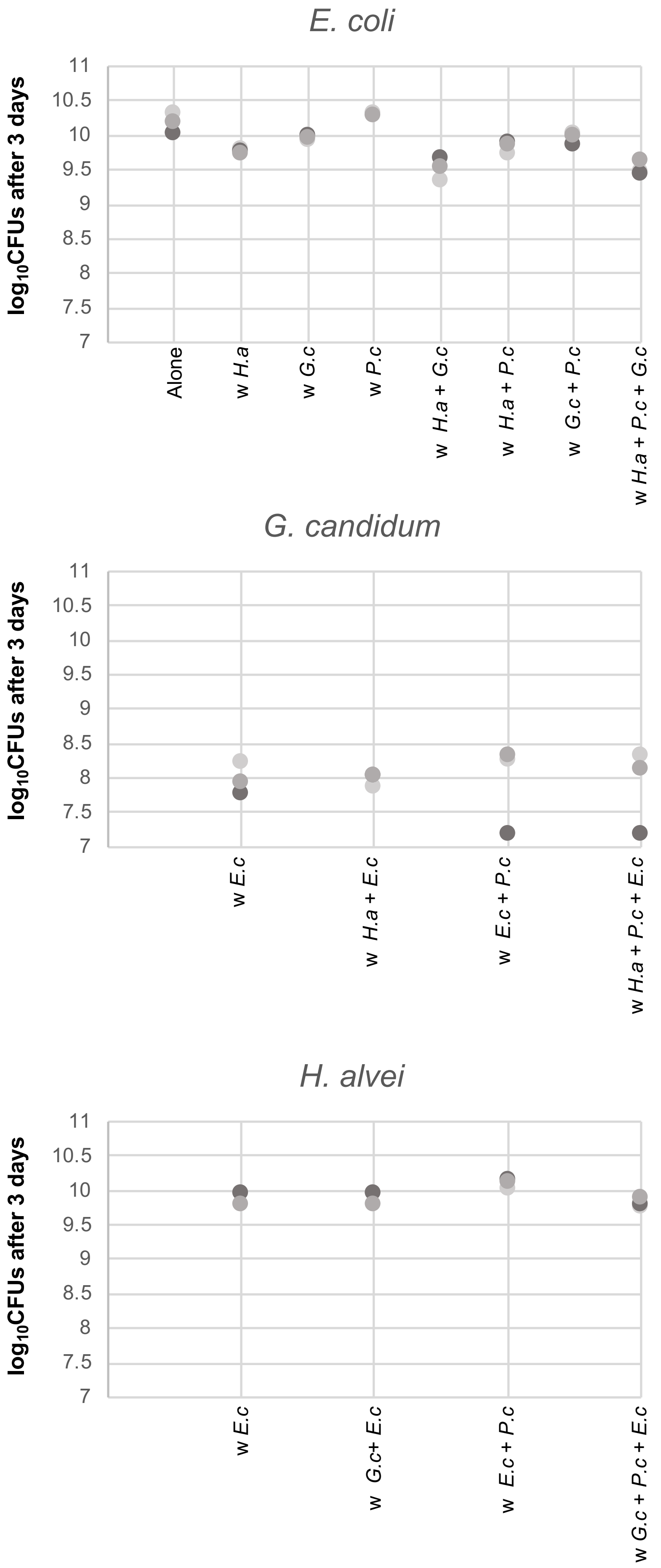


**Supplementary Figure 1: Growth of the community species in the studied conditions.**
Here, we show the $\log_{10}$ of CFUs for each species in the different conditions for each replicate after 3 days of growth.
Dark gray: Replicate 1; Medium gray: Replicate 2; Light grey: Replicate 3
*E.c: E. coli; H.a: H. alvei; G.c: G. candidum; P.c: P. camemberti*

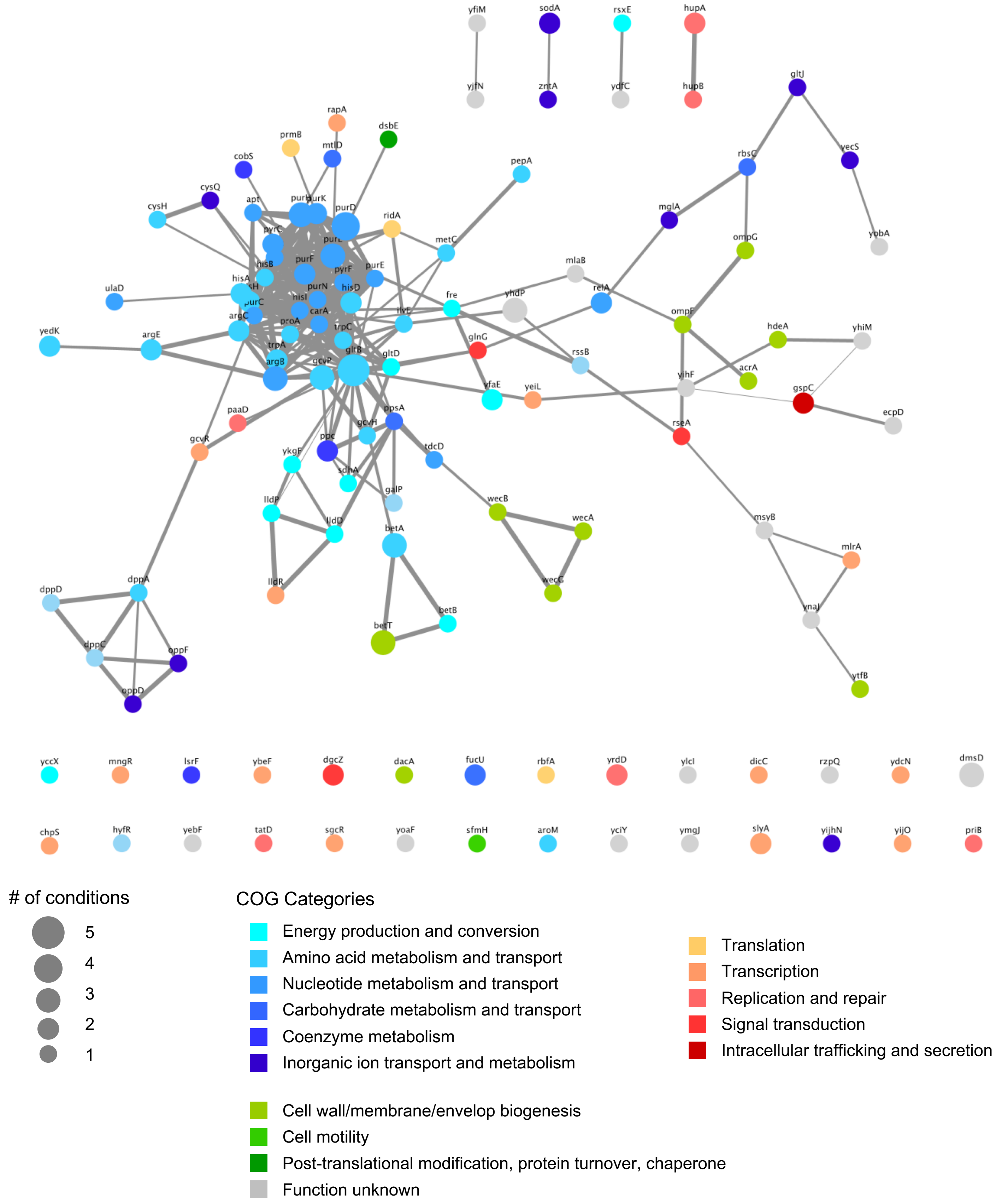


**Supplementary figure 2: Functional analysis of interaction-associated genes with negative IFEs.**
STRING network of the genes associated with negative IFEs (Nodes). Edges connecting the genes represent both functional and physical protein association and the thickness of the edges indicates the strength of data support (minimum required interaction score: 0.4 – medium confidence). Nodes are colored based on their COG annotation and the size of each node is proportional to the number of interactive conditions in which that given gene has been found associated with a significant IFE.

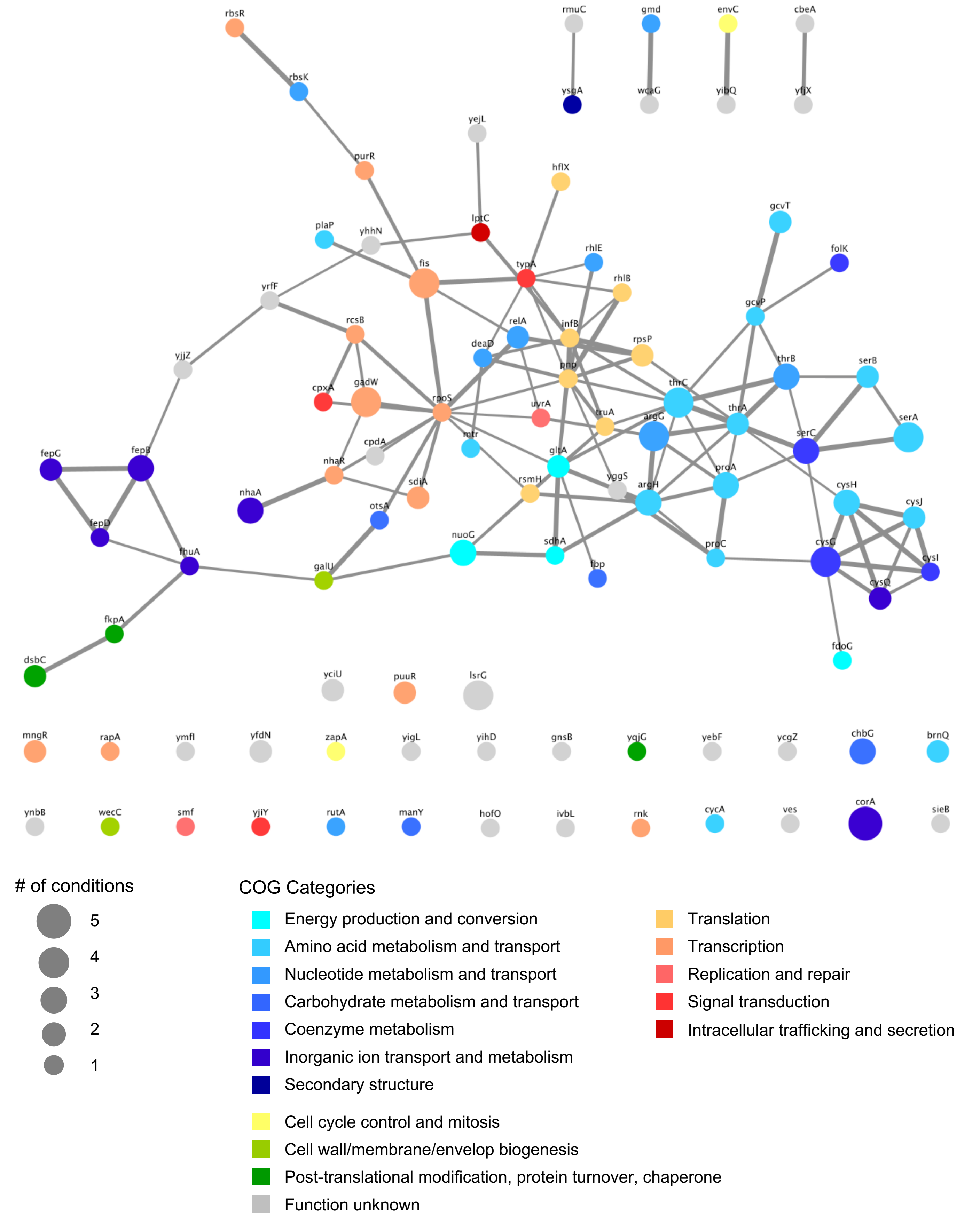


**Supplementary figure 3: Functional analysis of interaction-associated genes with positive IFEs.** STRING **n**etwork of the genes associated with positive IFEs (Nodes). Edges connecting the gens represent both functional and physical protein association and the thickness of the edges indicates the strength of data support (minimum required interaction score: 0.4 – medium confidence). Nodes are colored based on their COG annotation and the size of each node is proportional to the number of interactive conditions in which that given gene has been found associated with a significant IFE.

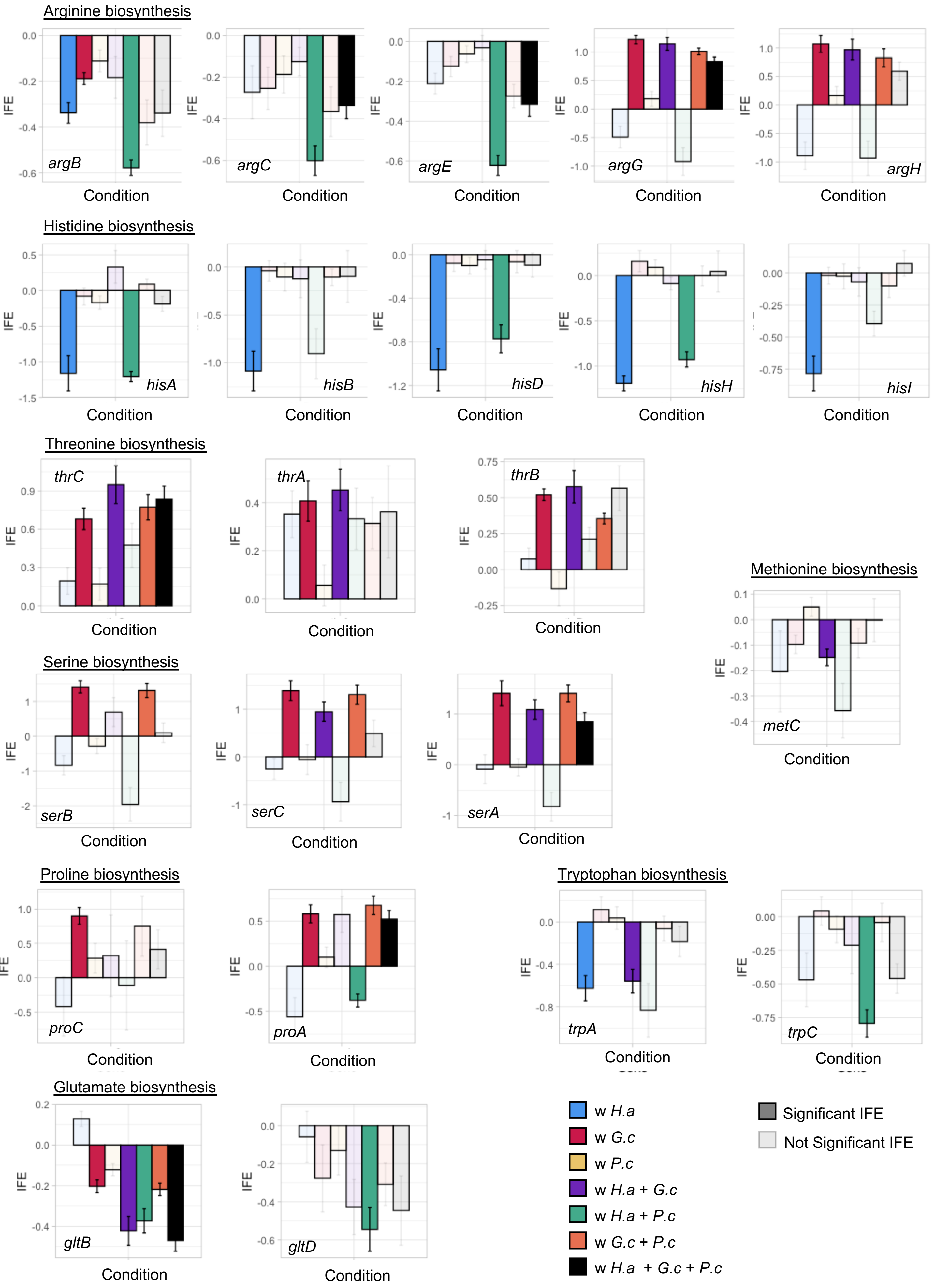


**Supplementary figure 4: IFE profiles of Amino acid biosynthesis genes identified in this study**

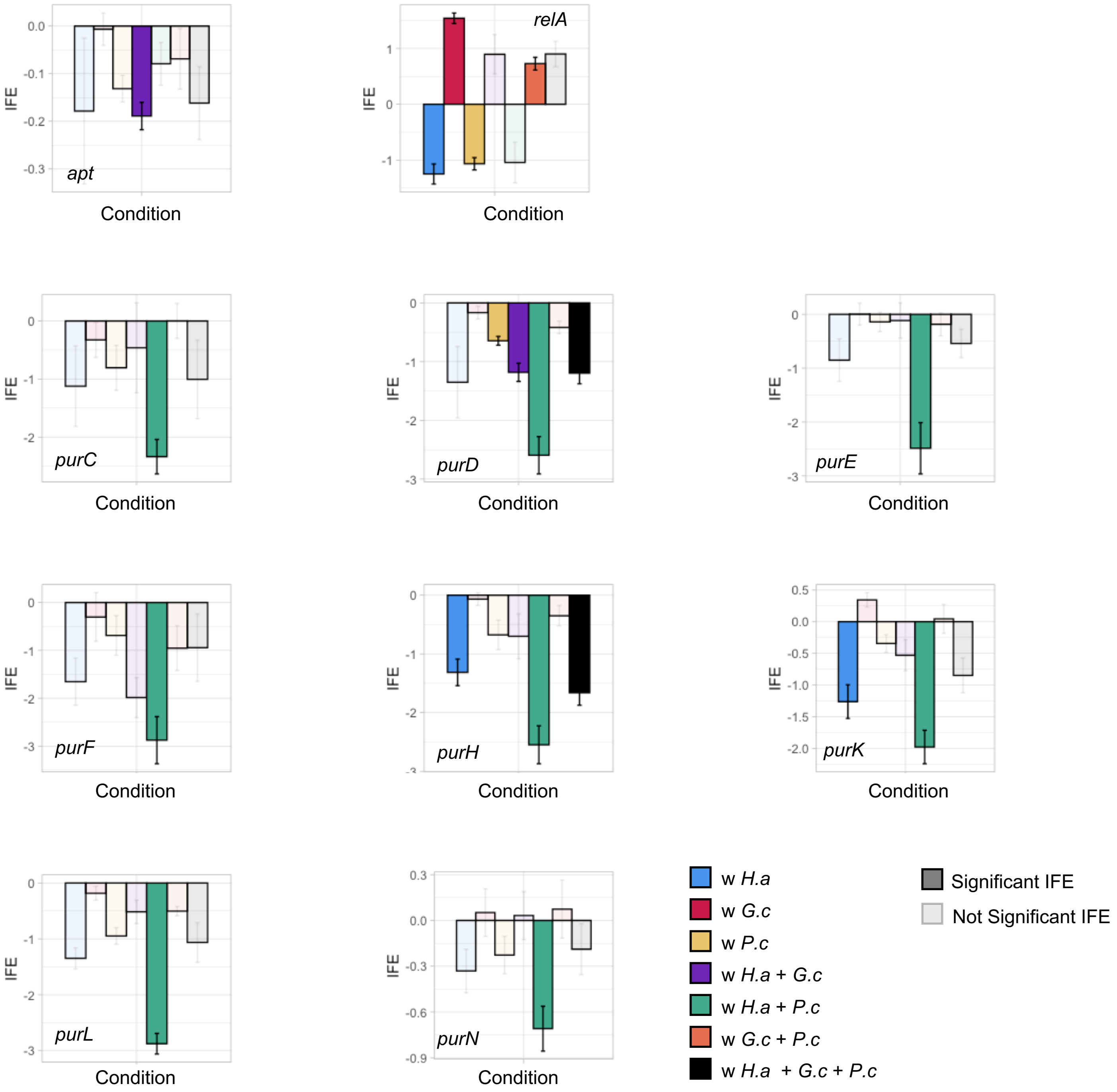


**<u>Supplementary figure 5: IFE profiles of Purine biosynthesis associated genes identified in this study</u>**

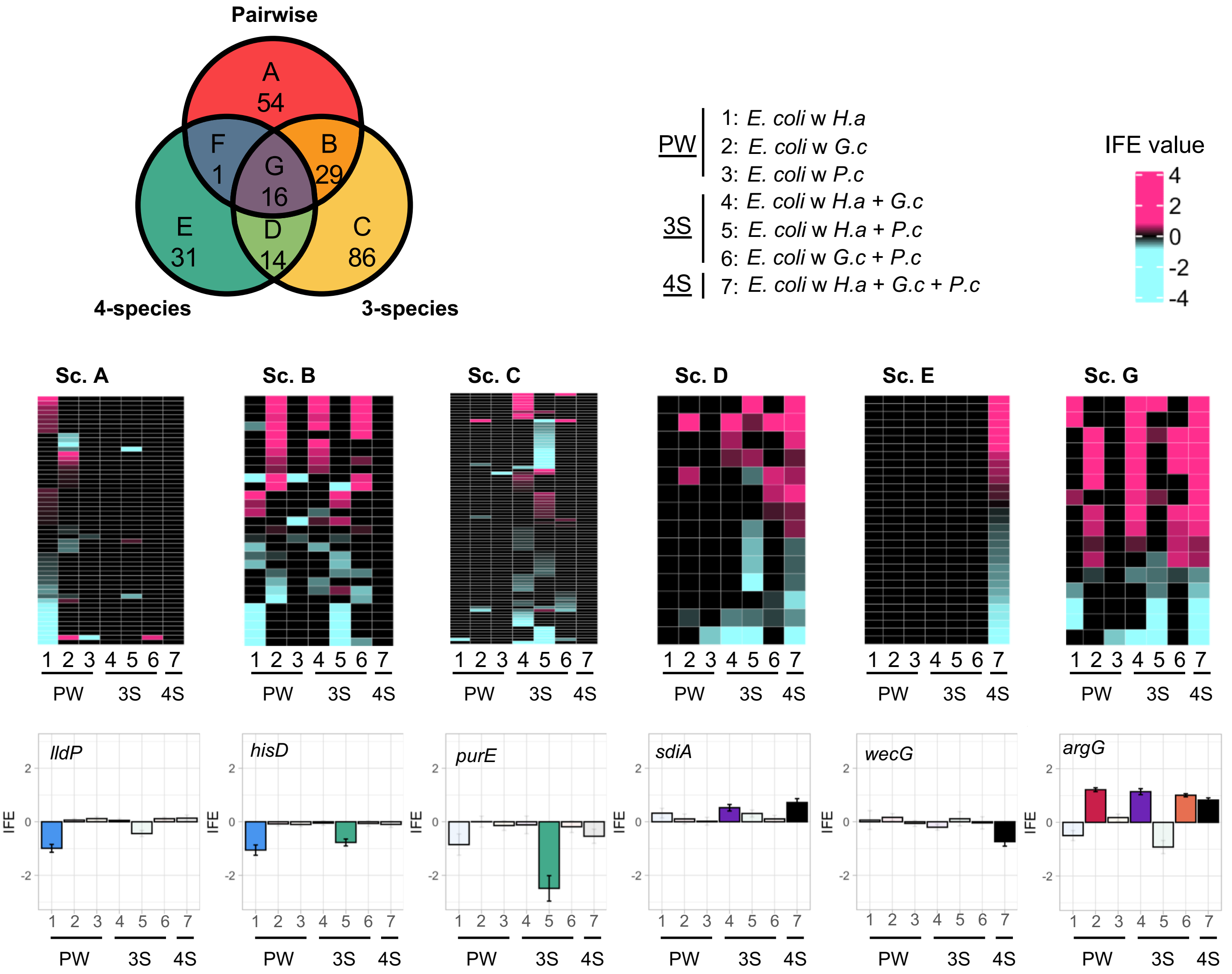


**Supplementary Figure 6: Comparison of interaction-associated genes across the different levels of community complexity.**
The Venn Diagram identifies 7 different possible scenarios (A to G) of interaction-associated genes conservation. Each scenario is then illustrated by a heatmap of the corresponding genes IFE values in all conditions (only the significant IFE are shown) along with the IFE profile of one example gene found in the scenario (Significant IFE: plain color, non-significant IFE: transparent color).

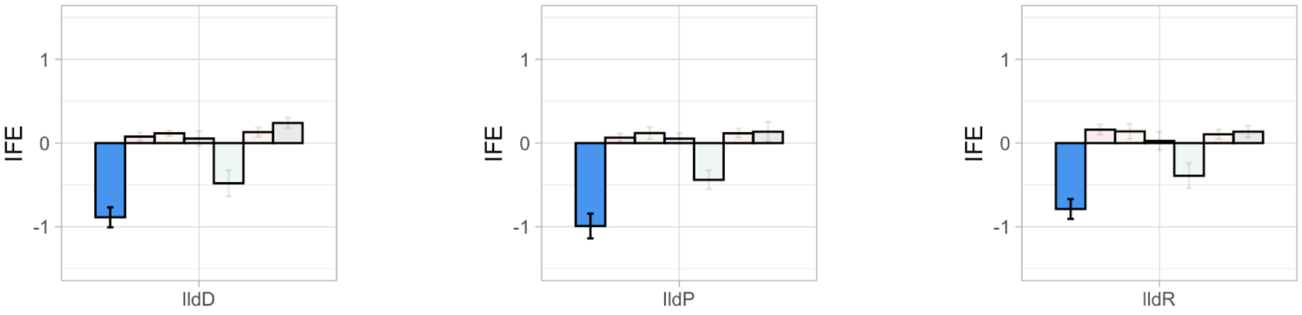


**Supplementary figure 7: IFE profiles of lactate metabolism genes.**

Maintained and Dropped 2-species interaction-associated gene in 3-species cultures

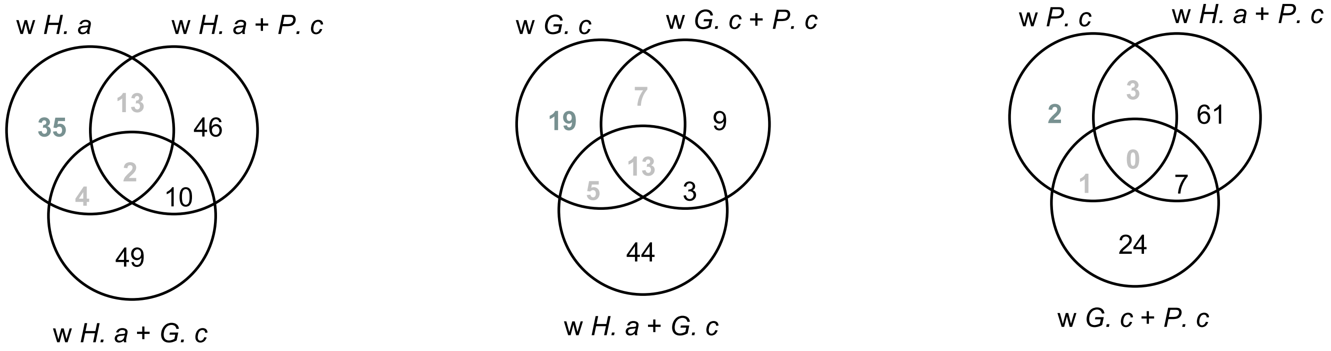


Emerging interaction-associated genes and 2-species maintained interaction-associated genes in 3-species cultures

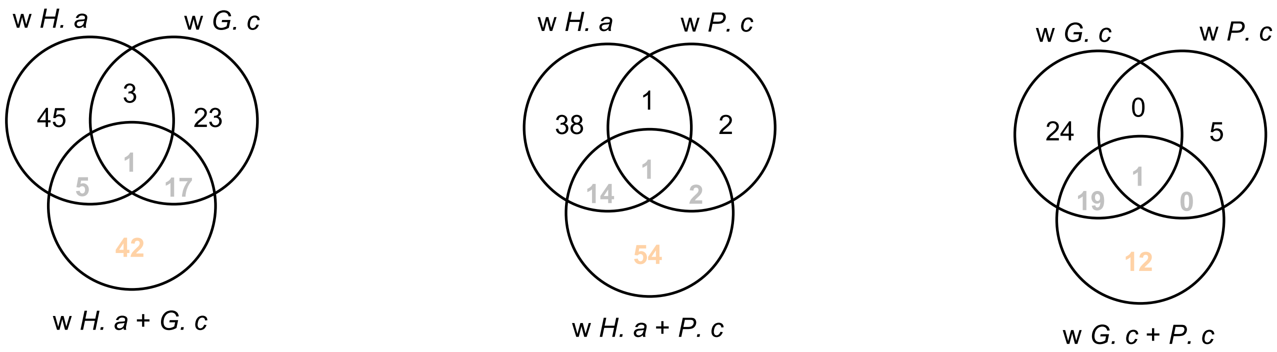


*H. a*: *Hafnia alvei*
*G. c*: *Geotrichum candidum*
*P. c*: *Penicillium camemberti*

**Supplementary figure 8: Condition specific comparison of interaction-associated genes for 2 and 3-species conditions**

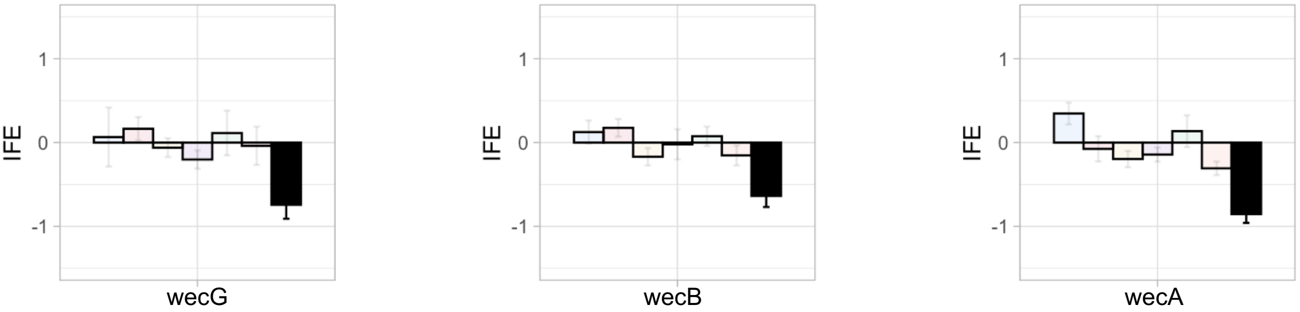


**Supplementary figure 9: IFE profiles of Enterobacterial Common Antigen (EAC) genes**

3-species dropped interaction-associated genes and 3-species maintained interaction-associated genes in 4-species cultures

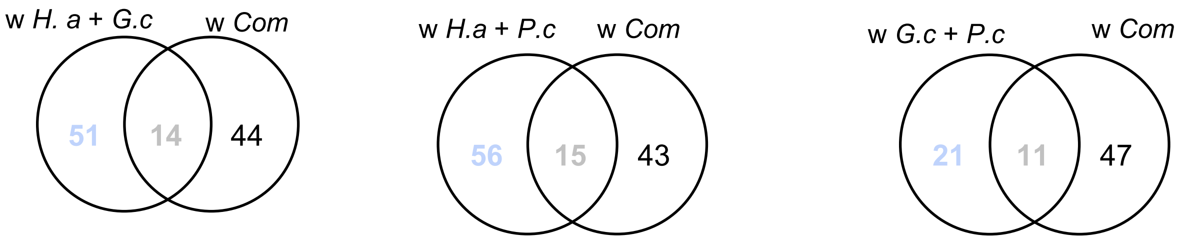


*H. a*: *Hafnia alvei*
*G. c*: *Geotrichum candidum*
*P. c*: *Penicillium camemberti*
*w Com= Hafnia alvei + Geotrichum candidum + Penicillium camemberti*

**Supplementary figure 10: Condition specific comparison of interaction-associated genes for 3 and 4-species condition**

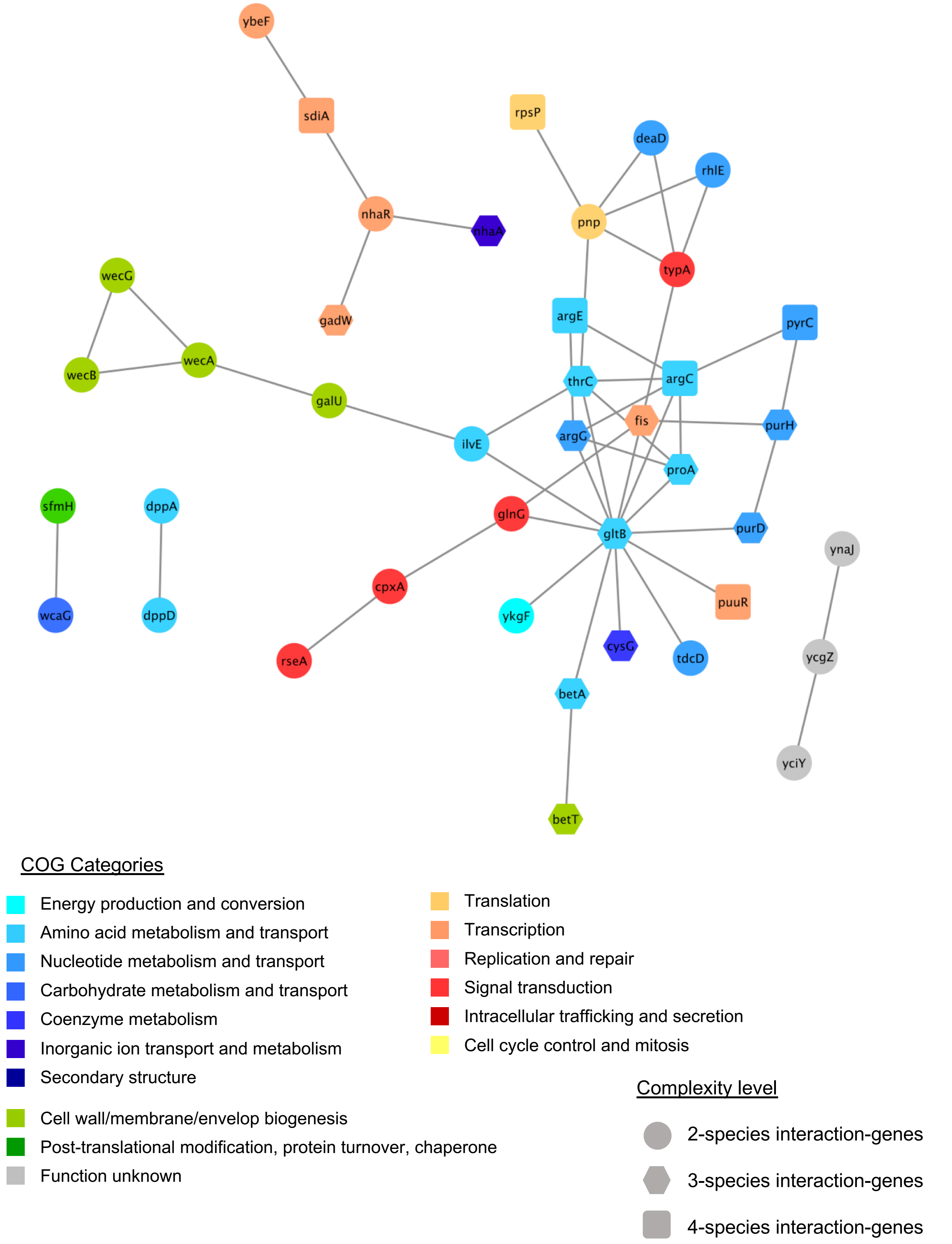


**Supplementary figure 11: Functional network of the 4-species interaction-genes and their origin**
STRING network of the genes (Nodes) associated with interactions in the 4-species condition. Edges connecting the genes represent both functional and physical protein association and the thickness of the edges indicates the strength of data support (minimum required interaction score: 0.4 – medium confidence). Nodes are colored based on their COG annotation and the shape of each node represents the level of community complexity the 4-species interaction-genes originate from.

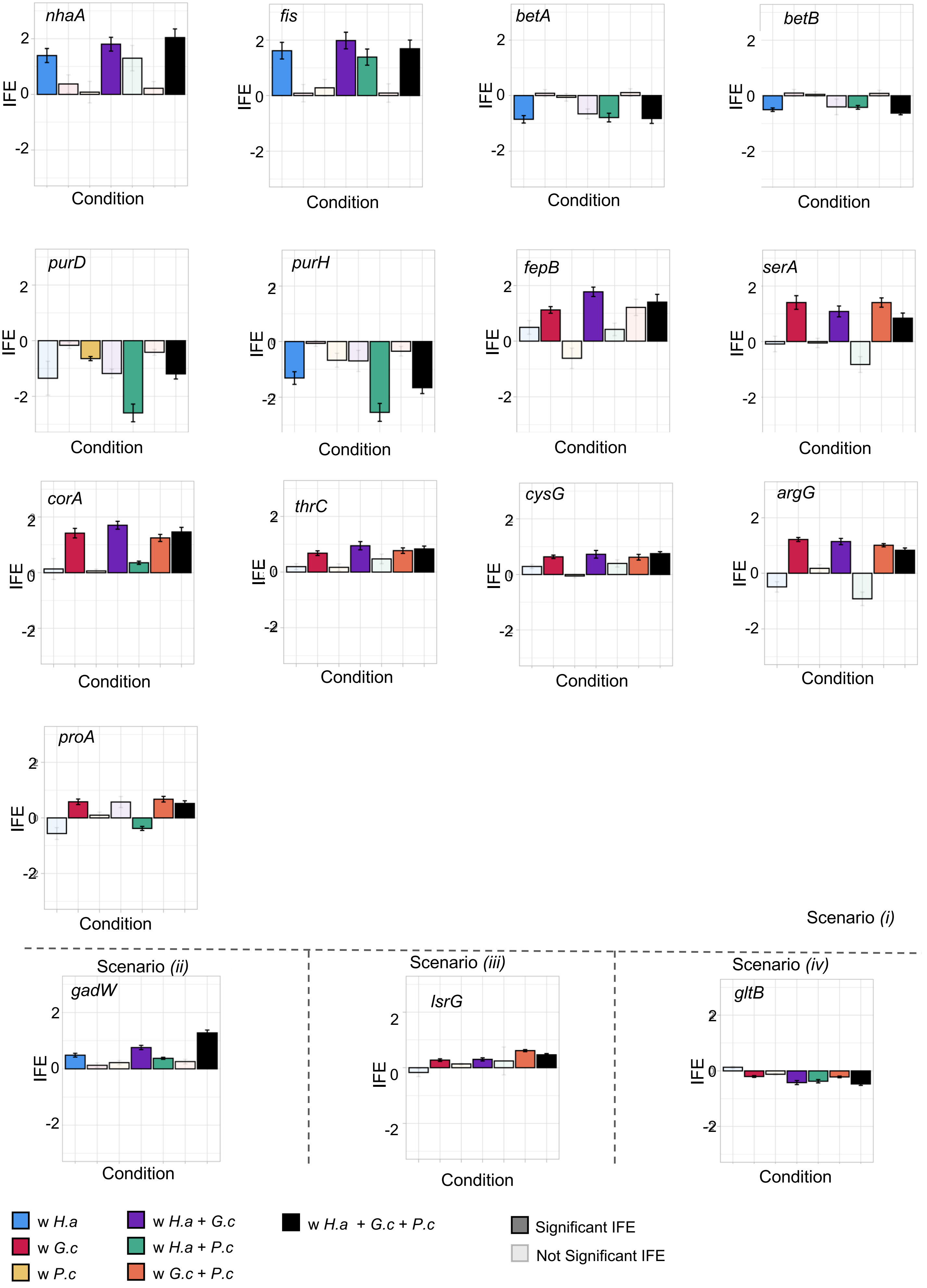


**Supplementary figure 12: IFE profiles of the 16 2-species interaction genes maintained up to 4-species**

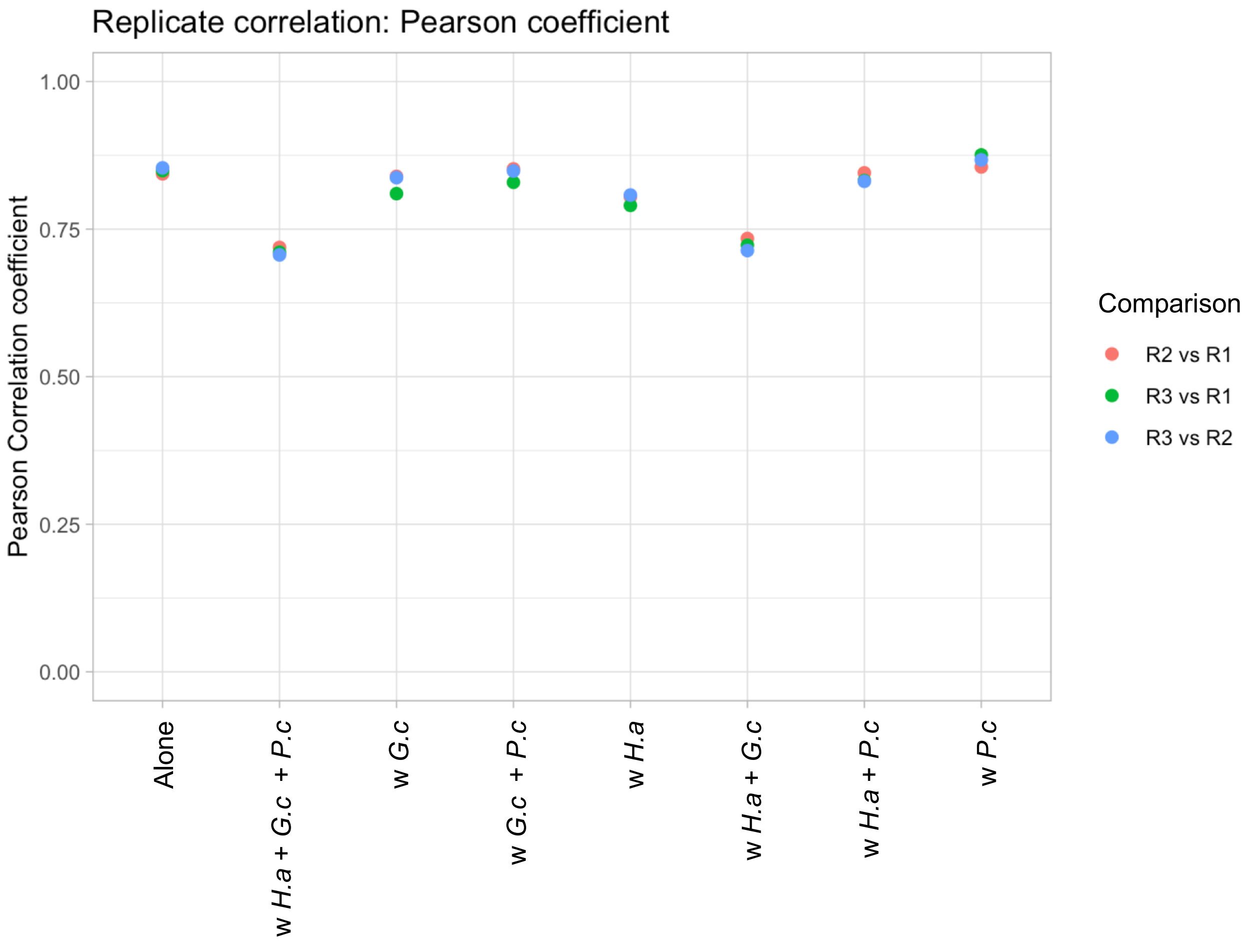


**Supplementary figure 13: Pearson correlation of gene fitness across replicates**

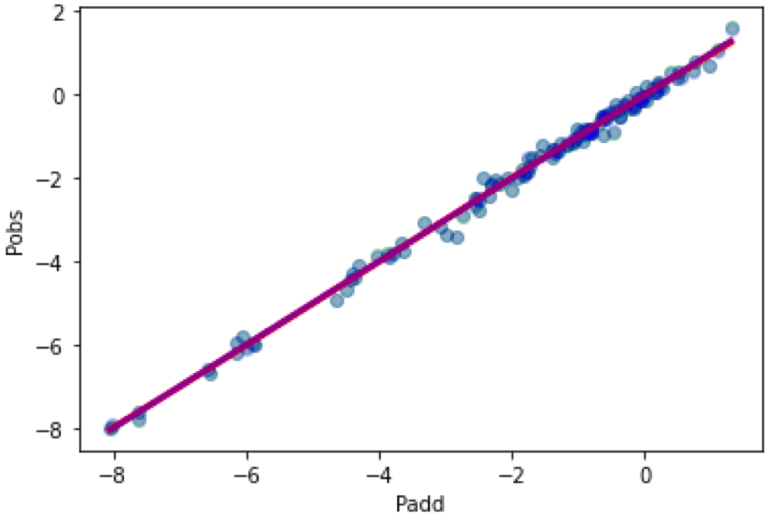


**<u>Supplementary figure 14: Non-linearity analysis of IFE in the Epistatis model</u>**
Predicted IFEs from an additive model (Padd) plotted against the Observed IFEs (Pobs) for the 16 genes associated with interactions from 2-species up to 4-species condition. No deviation from the identity line indicate that the IFE combine linearly and that there is no need for non-linearity correction (Sailer and Harms 2017).
